# An H3K9me3^high^ epigenetic state constrains neuronal activation and synaptic remodeling in the aged dentate gyrus

**DOI:** 10.64898/2026.11.13.766582

**Authors:** Yunuen Moreno-López, Carlos J. Rodriguez-Ortiz, Antonio Fuentes-Ibañez, Citlali A. Suárez-Rangel, Christopher W. Butler, Nicole C. Berchtold, G. Aleph Prieto

## Abstract

How epigenetic states shape neuronal responses to experience and memory remains poorly understood. Here, using a quantitative, single-event approach, our study identifies a subset of neurons that accumulate H3K9me3 (H3K9me3^high^ neurons) via the methyltransferase Suv39h1, specifically in the dorsal dentate gyrus (dDG) of aged mice. FACS coupled to RNA-seq showed that dDG-H3K9me3^high^ neurons express a transcriptional state that fails to induce plasticity machinery despite engaging memory-stabilization programs. Using our novel NucleuSynapse-Tag approach, we found that H3K9me3 accumulation interferes with neuronal activation and synaptic remodeling in response to behavioral stimulation in aged mice. Further, genetically inhibiting Suv39h1 to lower dDG-H3K9me3^high^ numbers improved memory in aged mice. Together, these findings reveal a region- and cell-type-specific epigenetic mechanism linking impaired neuronal responsiveness to age-related memory decline, providing mechanistic insight into the therapeutic potential of targeting Suv39h1.

## Introduction

An accruing body of work indicates that epigenetic mechanisms, which generate dynamic and reversible changes to chromatin structure to modify gene accessibility to the basal transcription machinery, are integral to memory encoding and storage^1-4^. Furthermore, epigenetic dysregulation has been hypothesized to underlie cognitive decline with aging and neurodegenerative diseases such as Alzheimer’s disease (AD)^5-9^. Indeed, a recent study suggests that epigenetic differences in chromatin accessibility in the human hippocampus represent a more robust molecular signature of differences in cognitive aging than differential mRNA expression^10^. Neuroepigenetics thus represents a growing, promising new field not only for understanding mechanisms underlying cognitive function, but also for generating effective treatments to counteract cognitive decline.

Histone methylation has emerged as a highly dynamic epigenetic mechanism for the modulation of experience-dependent plasticity in mature animals^11,12^. Specifically, the trimethylation of histone H3 at lysine 9 (H3K9me3), which is involved in gene repression^13^ and heterochromatin formation^7,12^, plays a role in brain processes including emotional responsiveness^14,15^, stress^16,17^, and memory^18,19^. We^20^ and others^21,22^ have reported that increased H3K9me3 levels in the hippocampus are associated with memory deficits in aged mice. Consistently, pharmacological inhibition of H3K9me3 improves hippocampal-dependent memory in aged^20^ and AD^23^ transgenic mice, indicating that H3K9me3 negatively regulates memory in the context of aging and neurodegeneration. However, the cellular and molecular mechanisms linking H3K9me3 and memory are poorly characterized.

Previous work has demonstrated that neither the rate nor the baseline transcription are equal across neuronal cell types in the hippocampus^24^, suggesting that patterns of epigenetic marks, including H3K9me3, are cell-type-specific as well. We hypothesized that the age-related increase in H3K9me3 levels in the hippocampus^20,21^ reflects accumulation in neuronal subgroups and is region-specific (*i.e.*, it differentially accrues in CA1, CA3, and the dentate gyrus (DG)). To address this possibility, we quantified H3K9me3 levels using a cell-type-specific approach via flow cytometry. This analysis revealed that a subset of neurons expressing high H3K9me3 levels (H3K9me3^high^) emerges with age in the dorsal DG (dDG). We characterized the genetic fingerprint of H3K9me3^high^ neurons isolated by fluorescence-activated cell sorting (FACS) and, for the first time, tested whether H3K9me3 accumulation interferes with neuronal activation and synaptic protein remodeling. Finally, a battery of behavioral tasks was used to evaluate memory following the genetic inhibition of H3K9me3 in the dDG of aged mice. Overall, our data identify an epigenetic mechanism that negatively impacts gene expression, neuron activation, synaptic remodeling, and memory in aging.

## Results

### H3K9me3 accumulates with age in a subset of hippocampal neurons

To address whether neurons or glia mainly contribute to the increased H3K9me3 levels in the hippocampus of aged mice^20,21^, we used a highly quantitative approach based on nuclear fractionation from dissected hippocampi and flow cytometry analysis of thousands of nuclei^24^ (**Fig. 1a,b**). In mice, H3K9me3 levels were assessed at three ages: young (3–4 months old [m.o.]), middle-aged (13–15 m.o.) and aged (18 m.o.), as cognitive deficits are generally absent in youth but emerge in middle age and become evident in late life^25,26^. In nuclei identified by DAPI, NeuN immunostaining allowed distinguishing neuron (NeuN^+^) and non-neuron (NeuN^−^) subpopulations (**Fig. 1b**). Data show comparable numbers of DAPI^+^, NeuN^+^, and NeuN^−^ nuclei in young, middle-aged, and aged mice (**Fig. S1**), consistent with the notion of preservation of glia^27^ and neurons^27,28^ numbers in the aged hippocampus. We found detectable levels of H3K9me3 in both NeuN^+^ and NeuN^−^ populations across the lifespan; however, the H3K9me3 mark was present in more NeuN^+^ than NeuN^−^ nuclei (\**P* = 0.02, **Fig. 1c,e**). Analysis across the three ages revealed an age-related increase in H3K9me3 intensity in neurons (*P* = 0.004, 3 vs 15 m.o.; *P* = 0.004, 3 vs 18 m.o.) but not in NeuN^−^ cells (**Fig. 1c,f**). The broadening of the H3K9me3 signal intensity curve with aging suggests a non-homogeneous population; indeed, applying a cut-off value defining neuronal nuclei with highest levels of the H3K9me3 signal (H3K9me3^high^) revealed an age-dependent emergence of a NeuN^+^ H3K9me3^high^ subpopulation (*P* = 0.04, 3 vs 15 m.o.; *P* = 0.002, 3 vs 18 m.o.; **Fig. 1d,g**), an unexpected and exciting finding. Conversely, the percentage of neurons with undetectable H3K9me3 levels (H3K9me3^neg^) decreased with age (*P* = 0.04, 3 vs. 18 m.o.), with no changes in neurons with low levels (H3K9me3^low^) (**Fig. 1d,g**). As a critical parameter in our study, the cut-off value for defining the H3K9me3^high^ subpopulation by flow cytometry was rigorously set and applied to the full set of samples. The immunostaining protocol (permeabilization, antibody concentrations, incubation times) and flow cytometry settings (PMT voltages, gains, acquisition speeds) were kept constant. Using this strategy, we found that most hippocampal neurons contain detectable but heterogeneous H3K9me3 levels that increase with age, thus altering the proportion of H3K9me3^high^ relative to H3K9me3^low^ and H3K9me3^neg^ neurons in the mouse hippocampus.

**Fig. 1.**
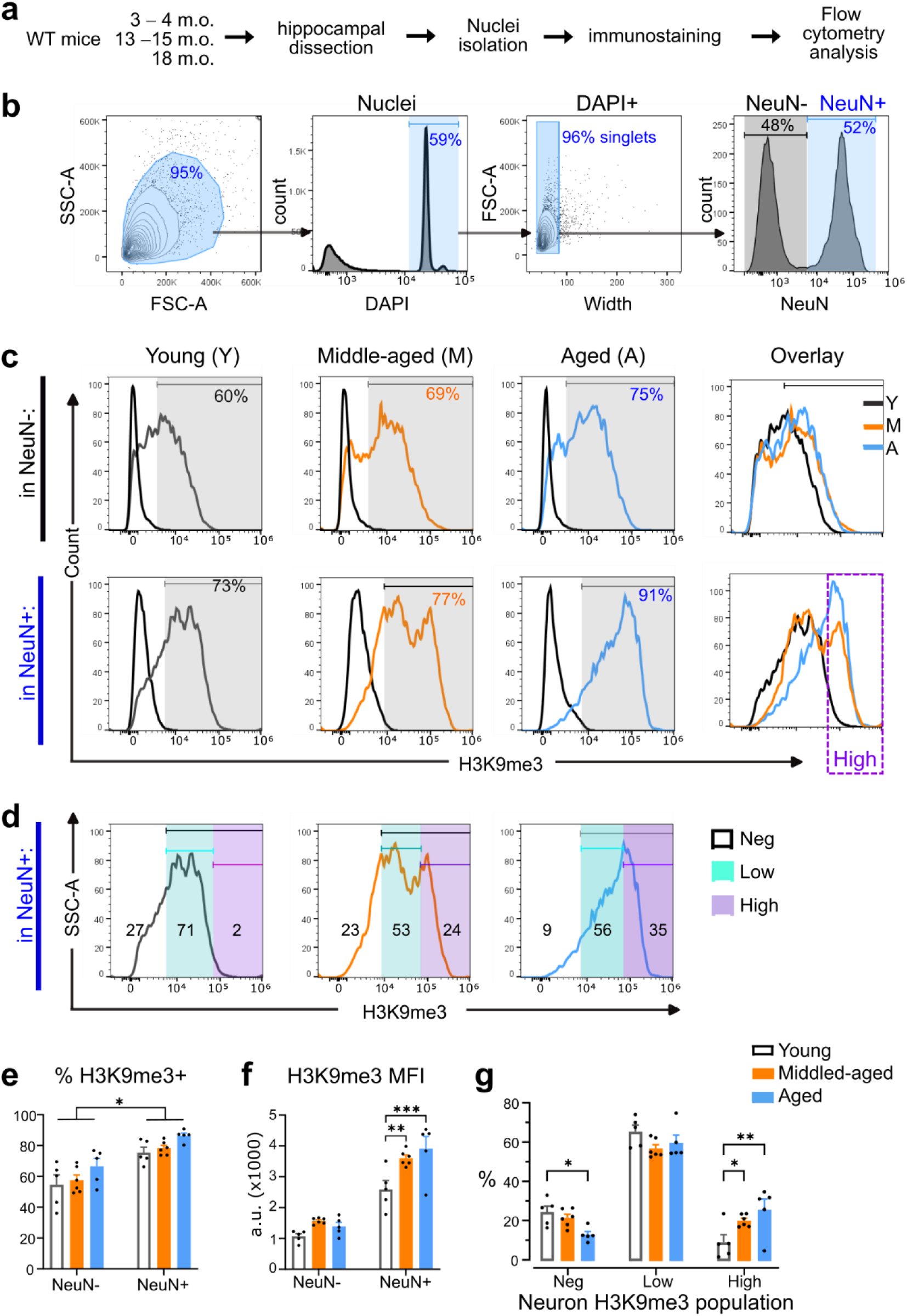
H3K9me3 accumulates in a subset of hippocampal neurons (H3K9me3^High^ neurons) with age. **a,** Workflow: WT mice at 3-, 15-, and 18-month-old (m.o.) were used for flow cytometry analysis of cell nuclei from hippocampi. **b,** Representative FACS plots show the hierarchical gating strategy to identify nuclei by size (FSC-SSC plot) and DAPI staining and further distinguish glia and neurons by NeuN labeling in singlets (events selected by doublet discrimination). **c,** H3K9me3 levels were assessed in young (3 m.o., gray trace), middle-aged (15 m.o., orange trace), and aged (18 m.o., blue trace) mice; each histogram shows the FMO control as a black trace. Histogram overlay identified neurons with high H3K9me3 signal (violet square). **d,** By establishing a cut-off value defining nuclei with the highest levels of H3K9me3, it was possible to study the abundance of three populations of neurons with undetectable (H3K9me3^Neg^), low (H3K9me3^Low^), and high levels of H3K9me3 (H3K9me3^High^) across age in the hippocampus. **e,** H3K9me3 labeling was more frequent in NeuN+, relative to NeuN-nuclei, *P* < 0.0001, main effect of “age”, \**P* = 0.01, Two-way ANOVA effect of cell population. **f,** Mean fluorescence intensity (MFI) of H3K9me3 increased with age in neurons only; \**P* = 0.0006, Two-way ANOVA effect of age, Sidak’s *post hoc*: \*\**P* = 0.004, \*\*\**P* = 0.0004. **g,** Bar graph shows the percentage of H3K9me3^Neg^, H3K9me3^Low^ and H3K9me3^High^ hippocampal neurons in young (n = 5), middle-aged (n = 6) and aged (n = 5) mice. \**P* < 0.0001, Two-way ANOVA main effect of H3K9me3-defined population, Sidak’s *post hoc*: \**P* = 0.04, \*\**P* = 0.002. Data are represented as mean ± s.e.m.

### Pharmacological inhibition of Suv39h1 reduces H3K9me3^high^ numbers

In aged and AD mice, hippocampus-dependent memory can be restored by reducing H3K9me3 via ETP69^20,23^, a small-molecule inhibitor of Suv39h1 (the HMT that trimethylates H3K9^29^). Using flow cytometry, we investigated how ETP69 affects the distribution of the H3K9me3-defined neuron subpopulations. Middle-aged (15 m.o.) male mice were administered ETP69 (10 mg/Kg, i.p.) or vehicle, and hippocampal tissue was collected after 24 h. ETP69 selectively lowered the H3K9me3^high^ subpopulation, whereas increased H3K9me3^low^ neurons (*P* = 0.007) (**Fig. 2a,b**). Consequently, the H3K9me3^low^/H3K9me3^high^ ratio increased in ETP69-treated animals (*P* = 0.01; **Fig. 2c**), thereby shifting the balance of the H3K9me3-defined neuron populations towards the profile of a younger brain. Taken together with our previous finding that ETP69 restores cognition in aged mice^20^, these data suggest that the selective reduction of the H3K9me3^high^ population enables the memory-enhancing effect of ETP69. Next, we focused on identifying hippocampal regions vulnerable to H3K9me3 accumulation with age.

**Fig. 2.**
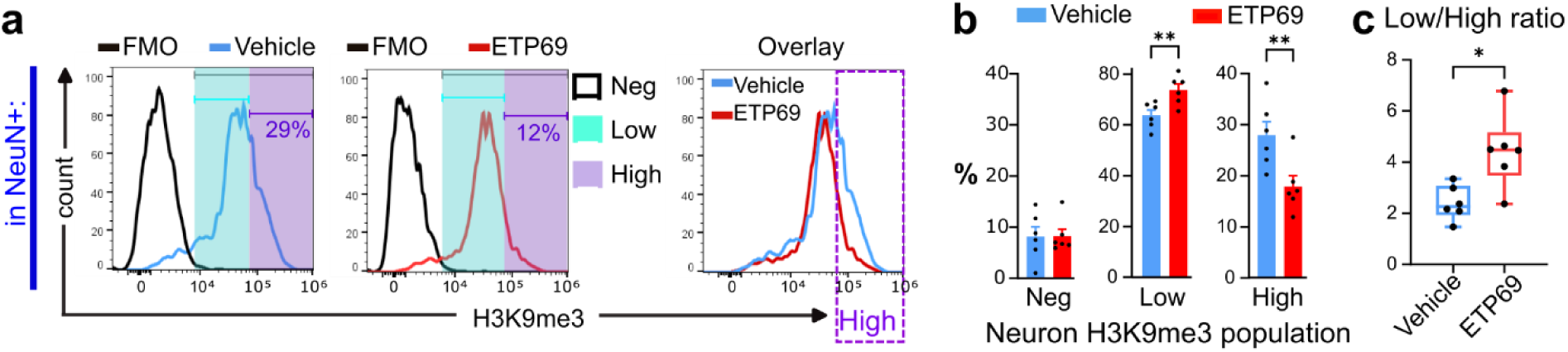
Pharmacological inhibition of SUV39H1 by ETP69 reduces the proportion of H3K9me3^High^ neurons in the aged hippocampus. Dissected hippocampi from mice injected with ETP69 1 mg/kg (i.p.) or vehicle were used for isolating cell nuclei, which were analyzed by flow cytometry. **a,** Neuronal nuclei were identified by size, DAPI and NeuN staining. To quantify the abundance of neurons with undetectable low (H3K9me3^Low^) and high (H3K9me3^High^) levels of H3K9me3, we use the cut-off value defining nuclei with highest levels of H3K9me3 signal (*see* **Fig. 1**). **b,** Bar graphs show the percentage of H3K9me3^Neg^, H3K9me3^Low^ and H3K9me3^High^ hippocampal neurons in vehicle (n = 6) and ETP69 (n = 6) groups of mice, \**P* < 0.0001, Two-way ANOVA main effect of H3K9me3-defined population, Sidak’s *post hoc*: \*\**P* = 0.007. **c,** H3K9me3^Low^/H3K9me3^High^ (Low/High) ratio was compared in vehicle and ETP conditions. \**P* = 0.01, two-tailed unpaired *t*-test. Data are represented as mean ± s.e.m.

### H3K9me3^high^ neurons emerge with age selectively in the dorsal DG

We investigated the distribution of the critical subpopulation of H3K9me3^high^ neurons to determine if certain hippocampal subregions were preferentially affected. To this end, we identified neurons of major hippocampal subfields (CA1, CA3, and DG) by flow cytometry using a specific antibody panel (NeuN, Prox1, Ctip2) in isolated nuclei (DAPI^+^), as previously reported.^24^ Four neuronal subpopulations were identified: DG (Prox1^+^Ctip2^+^), CA1 (Prox1^−^Ctip2^+^), and CA3 (Prox1^−^Ctip2^−^) excitatory neurons, and vasoactive intestinal polypeptide-expressing interneurons (iVIP) (Prox1^+^Ctip2^−^) (**Fig. 3a-c**). Data show that the relative neuron abundance of CA1, CA3, DG, and iVIP neurons is comparable between young (3–4 m.o.) and aged (18 m.o.) mice (**Fig. 3d**). Notably, we found that the H3K9me3^high^ subpopulation emerges with age selectively in the DG (*P* < 0.0001, **Fig. 3e,h**). Supporting the idea that DG neurons are more prone to H3K9me3 accumulation, we found that the abundance of H3K9me3^neg^ neurons was the lowest in DG compared to all other populations in both young and aged mice (*P* < 0.0001, **Fig. 3e, f**). These data show that the age-related increase of H3K9me3 levels is region-specific and differentially affects the CA1, CA3, and DG, with the latter selectively increasing the number of H3K9me3^high^ neurons, as assessed in the whole hippocampus.

**Fig. 3.**
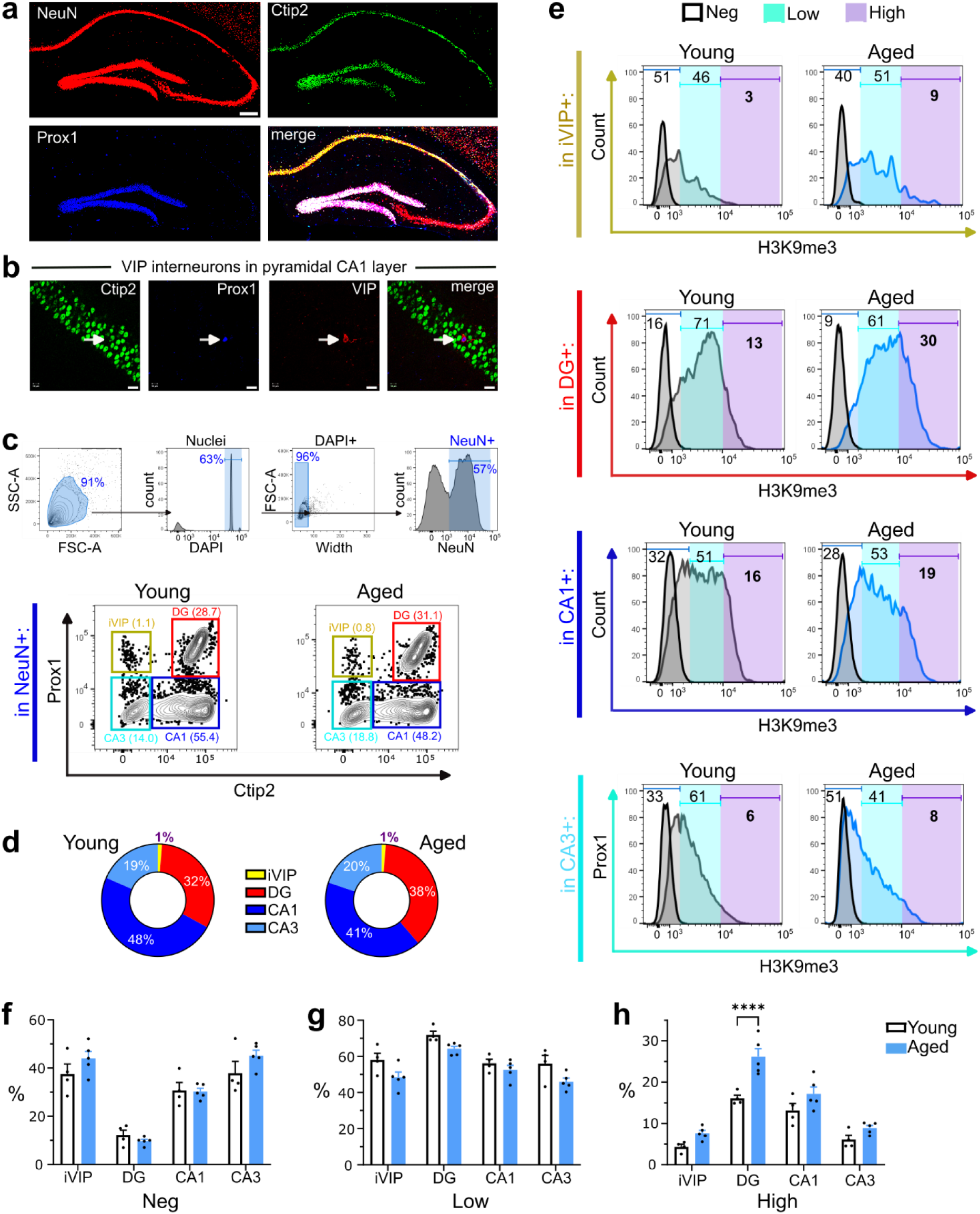
H3K9me3^High^ neurons selectively emerge in DG of aging mice. **a,** Confocal imaging of NeuN, Prox1, and Ctip2 proteins in the hippocampus. Scale bar= 200 μm. **b,** Images confirming that VIP^+^ interneurons can be identified as Ctip2^Neg^Prox1^+^ cells, as shown in the pyramidal CA1 layer. Scale bar= 20 μm. **c,** The whole hippocampus was dissected to isolate cell nuclei for hierarchical flow cytometry analysis to identify NeuN^+^ events (illustrated by arrows). For identifying hippocampal regions, a specific antibody panel (NeuN, Prox1, Ctip2) combined with DAPI identified DG (Prox1^+^Ctip2^+^), CA1 (Prox1^−^Ctip2^+^) and CA3 (Prox1^−^Ctip2^−^) excitatory neurons, and VIP interneurons (iVIP) (Prox1^+^Ctip2^−^). Representative density plots show iVIP, DG, CA1, and CA3 neuron populations in young (4 m.o.) and aged (18 m.o.) mice; % are shown in parentheses. **d,** Circle charts show the proportion of iVIP, DG, CA1, and CA3 neurons in young and aged mice. **e,** Representative histograms of H3K9me3 levels, and H3K9me3-defined populations, in iVIP, DG, CA1, and CA3 neurons in young and aged mice. (**f-h),** Percentage of (**f**) H3K9me3^Neg^, (**g**) H3K9me3^Low^, and (**h**) H3K9me3^High^ neurons in iVIP, DG, CA1, and CA3 neurons, comparing young (n = 4) and aged (n = 5) mice. Two-way ANOVA, main effects of age and hippocampal subfield (*P* < 0.0001), Sídák’s *post hoc* analysis: \*\*\*\**P* < 0.0001. Data are represented as mean ± s.e.m.

Next, we tested potential differences in the number of DG H3K9me3^high^ neurons by comparing dorsal and ventral hippocampal regions, which are associated with cognitive and emotional processes, respectively^30^. In 18-month-old mice, DG neurons were more abundant in the dorsal hippocampus (40.7 % vs 16.8 % neurons, dorsal vs ventral, *P* < 0.0001; **Fig. 4a,b**). Consequently, the absolute number of DG H3K9me3^high^ neurons was higher in the dorsal hippocampus (*P* = 0.004, dorsal vs ventral; **Fig. 4e**), even though the percentages of H3K9me3^high^ neurons were comparable in dorsal and ventral regions (**Fig. 4c,d**). These results indicate that hippocampal H3K9me3^high^ neurons become abundant with age selectively in the dDG, with no apparent differences between left and right hippocampi (**Fig. S2**).

**Fig. 4.**
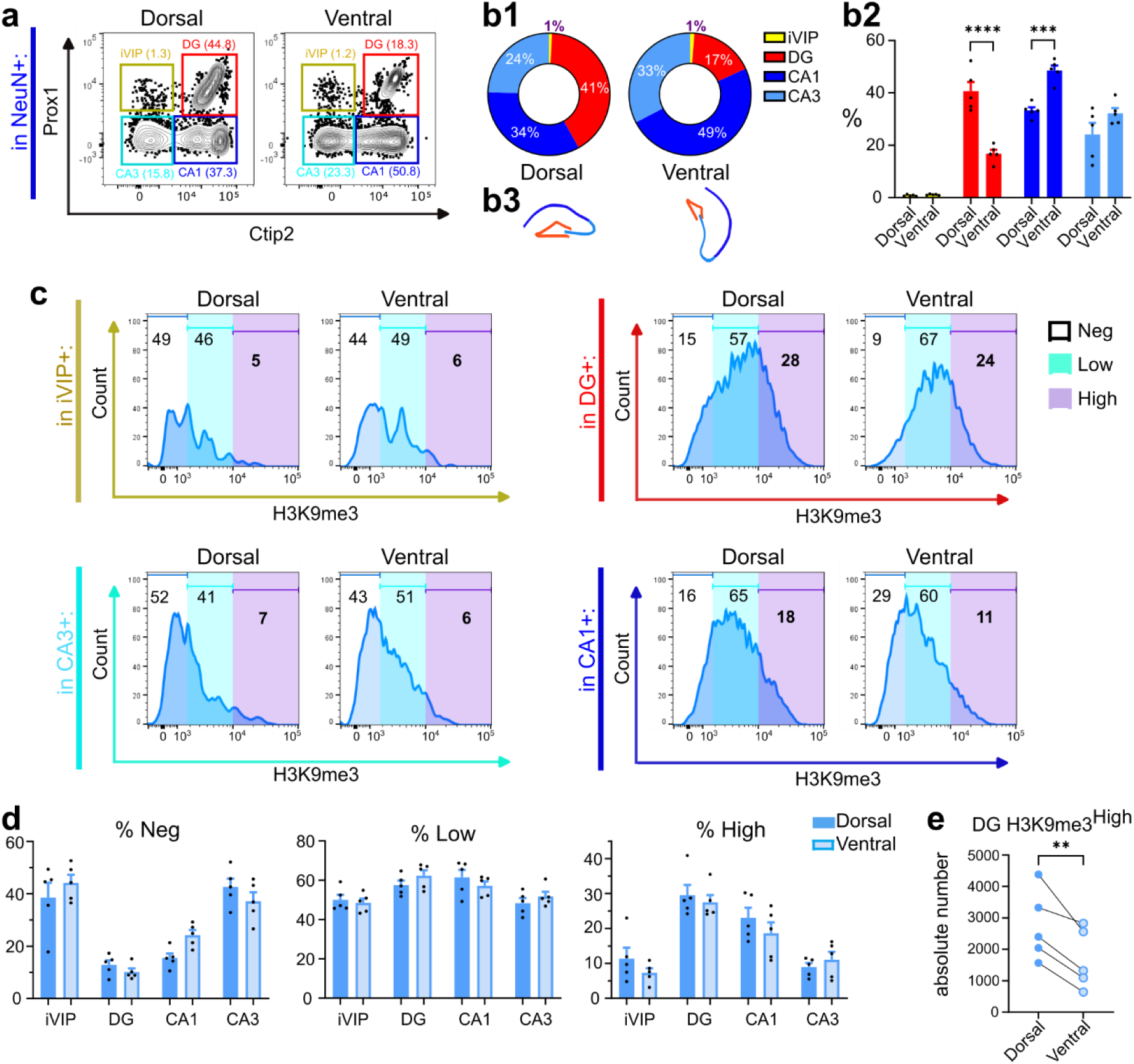
H3K9me3^High^ neurons mainly accumulate in the *dorsal* region of the aged DG. **a,** Flow cytometry dissection of the hippocampus using Ctip2 and Prox1 markers in NeuN^+^ nuclei obtained from dissected dorsal and ventral regions. Representative FACS plots show iVIP, DG, CA1 and CA3 populations (% are shown in parentheses). **b,** Proportions (%) of iVIP, DG, CA1 and CA3 populations shown in circle charts (**b1**) and bar graphs (**b2**). Comparison of dorsal and ventral regions showed differential proportions of DG and CA1 neurons, consistent with anatomical differences between dorsal and ventral hippocampus (**b3**); Two-way ANOVA, main effect of region (*P* < 0.0001), Sídák’s *post hoc* analysis: \*\*\**P* = 0.002, \*\*\*\**P* < 0.0001 **c,** Representative histograms of H3K9me3 levels in iVIP, DG, CA1, and CA3 neurons in dorsal and ventral regions. **d,** Percentage of H3K9me3^Neg^, H3K9me3^Low^ and H3K9me3^High^ neurons in iVIP, DG, CA1, and CA3 subfields, comparing dorsal and ventral hippocampal regions in aged mice (n = 5). Two-way ANOVA, main effect of hippocampal subfield (P < 0.0001), with no effect for hippocampal dorsal and ventral regions (P > 0.05). **e,** Total counts of H3K9me3^High^ DG neurons in dorsal and ventral regions. \**P* = 0.008, two-tailed paired *t*-test. Data are represented as mean ± s.e.m.

### dDG-H3K9me3^high^ neurons engage engram programs but fail to recruit plasticity machinery upon learning

As H3K9me3 signals for gene repression^13^, we hypothesized that H3K9me3^high^ neurons fail to induce programs required for synaptic plasticity and memory consolidation. RNA-seq was used to profile FACS-sorted H3K9me3^high^ and H3K9me3^low^ dDG neurons from home-cage (HC) and contextual-fear-conditioned (CFC) mice (**Fig. 5a,b**), as CFC robustly induces long-term memory even in aged animals^9^. RNA-seq data were analyzed with PyDESeq2 under a nested paired design with BH-adjusted *P* < 0.1.

**Fig. 5.**
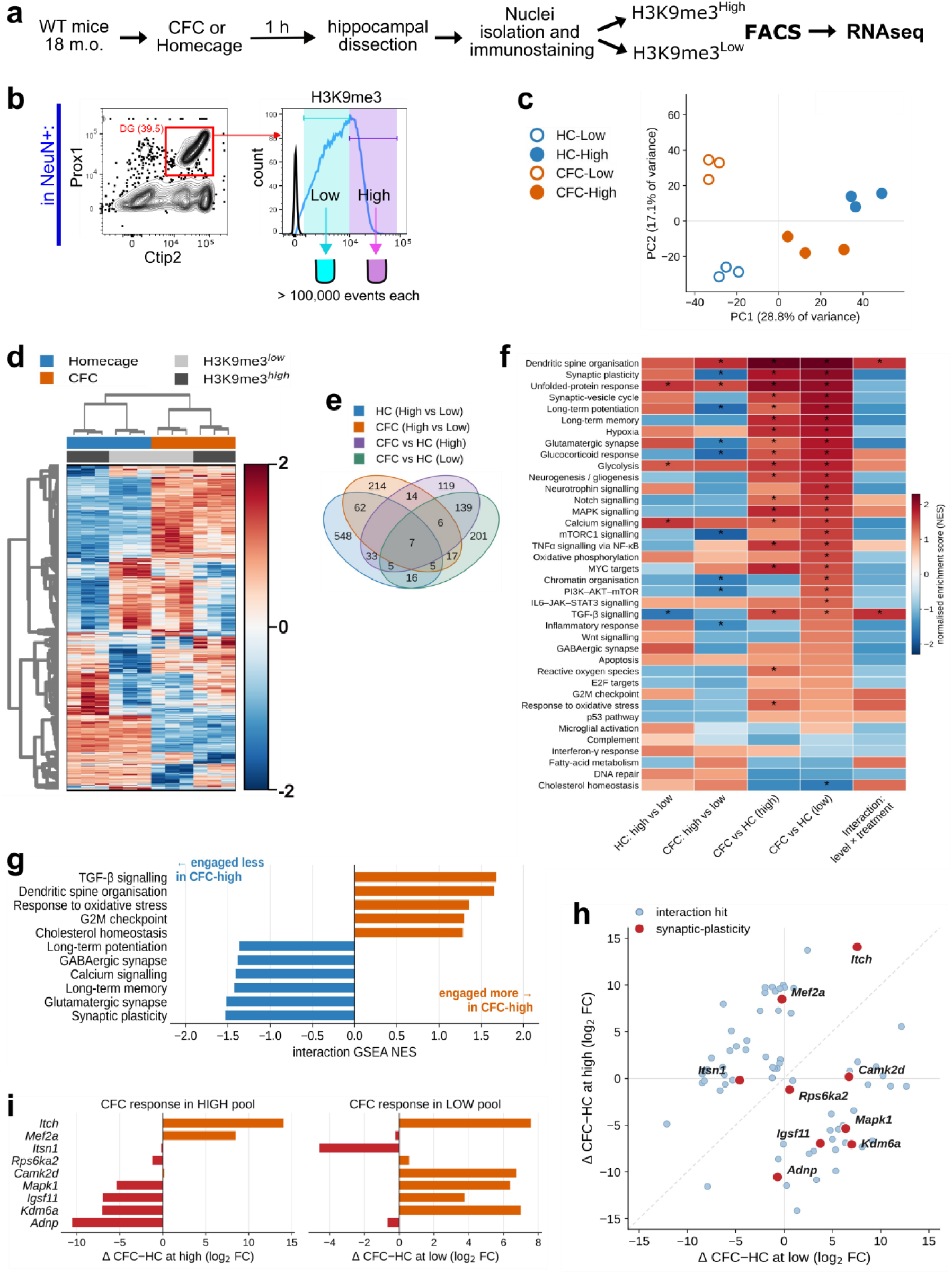
Dorsal-DG-H3K9me3^High^ neurons express a unique gene program that downregulates synaptic genes. **a**, Workflow: to induce neuron activation, 18-month-old (m.o.) mice were trained using the contextual fear conditioning (CFC) paradigm; as a control, we used a group of mice maintained in their habitual cage (homecage) (n=3 per condition). After 1 hour of CFC, dorsal hippocampi were dissected and processed for cell nuclei isolation and flow cytometry coupled to sorting (FACS), which allowed sorting of H3K9me3^Low^ and H3K9me3^High^ populations for RNA-seq. **b**, Representative cytometry plot (left) using Ctip2 and Prox1 markers in NeuN^+^ nuclei to identify DG neurons. The histogram (right) shows H3K9me3 levels in DG to select and sort H3K9me3^Low^ and H3K9me3^High^ populations; the FMO control is shown in black trace. **c**, Principal component analysis of the 12 RNA-seq samples, computed on the union of transcripts reaching P_adj_ < 0.10 in at least one of the four primary contrasts (n = 1,386 transcripts). PC1 (28.8 % of variance) separates H3K9me3^High^ from H3K9me3^Low^ samples — the within-animal axis — and PC2 (17.1 %) separates CFC from homecage samples. Each point is one sample; color denotes behavioral condition (homecage, blue; CFC, orange) and fill denotes H3K9me3 level (filled, high; open, low). Differential expression was tested at the transcript level with DESeq2 using a nested paired design, and P values were corrected within each contrast by the Benjamini–Hochberg false discovery rate; features with P_adj_ < 0.10 were considered significant. **d**, Clustered heatmap of the significant transcript set (n = 1,386 transcripts, Padj < 0.10 in at least one of the four pairwise contrasts from the nested paired model). Values are per-transcript z-scores of normalised log₂ expression after removal of within-condition animal effects for visualisation (group means preserved; all statistics were computed on unadjusted data). Transcripts (rows) and samples (columns) were ordered by unsupervised hierarchical clustering using correlation distance and average linkage. Annotation bars indicate condition (homecage, blue; CFC, orange) and H3K9me3 level (high, dark grey; low, light grey). **e**, Four-way Venn diagrams of significant features (P_adj_ < 0.10) at transcript level (left) and gene level (right), across the four primary contrasts: H3K9me3^High^ versus H3K9me3^Low^ in homecage animals; H3K9me3^High^ versus H3K9me3^Low^ in CFC animals; CFC versus homecage in the H3K9me3^High^ pool; and CFC versus homecage in the H3K9me3^Low^ pool. **f**, Pre-ranked gene set enrichment analysis across the curated 38-pathway library. Color is the normalized enrichment score (NES) for each pathway set (rows) in the four primary contrasts and the condition × H3K9me3-level interaction. Rows are the 27 pathway sets that reach FDR < 0.25 in at least one contrast, ordered by NES in the CFC versus homecage contrast within the H3K9me3^Low^ pool, from most positive at the top. Red, positive NES; blue, negative NES. Asterisks mark FDR < 0.25. **g**, Direction-resolved signed GSEA for the interaction contrast, showing the five most positive and six most negative pathway sets by interaction NES as determined by effect size rather than by significance. Negative values (blue) identify pathways engaged less in CFC-H3K9me3^High^ neurons — synaptic plasticity, glutamatergic synapse, long-term memory, calcium signaling, GABAergic synapse and long-term potentiation — and positive values (orange) identify pathways engaged more in that group: TGF-β signaling, dendritic spine organization, response to oxidative stress, G2M checkpoint and cholesterol homeostasis. **h**, Response of the 73 condition × H3K9me3-level interaction hits (Padj < 0.10) to conditioning in each pool. Each point is one interaction transcript, positioned by its CFC-versus-homecage log₂ fold change within the H3K9me3^low pool (x axis) and within the H3K9me3^high pool (y axis). The dashed diagonal marks identical responses in the two pools; points in the upper-left and lower-right quadrants respond in opposite directions. Red, the nine synaptic-plasticity genes among the hits (GOBP synaptic plasticity), labelled; blue, all other interaction hits. Kdm6a, Mapk1, Igsf11 and Rps6ka2 (lower right) are induced by conditioning in H3K9me3^low neurons but suppressed in H3K9me3^high neurons; Itch (upper right, off-scale marker) is induced in both pools. **i**, Gene-level dissection of the interaction for plasticity-machinery genes shown as the CFC response within each pool separately (H3K9me3^High^, left; H3K9me3^Low^, right); bars are Δ CFC − homecage log_2_ fold change, orange positive, red negative; note the independent x scales. Camk2d, Kdm6a, Mapk1 and Igsf11 are induced by CFC in the H3K9me3^Low^ pool but suppressed in the H3K9me3^High^ pool, with Rps6ka2 showing the same reversal more weakly. Itch is induced in both pools and more strongly in the H3K9me3^High^ pool, Mef2a is induced only in the H3K9me3^High^ pool, and Adnp and Itsn1 are suppressed in both, with greater suppression of Adnp in the H3K9me3^High^ pool.

Principal component analysis of significant transcripts across the four primary contrasts (n = 1,386 transcripts; 1,167 genes) separated CFC from HC samples with the H3K9me3 axis (PC1 (28.8%)) accounting for more variance than condition PC2 (17.1%) (**Fig. 5c**). Pseudogenes, lncRNAs, and small ncRNAs (biotypes canonically silenced by H3K9me3) were less expressed in the H3K9me3^high^ pool (**Fig. S3a**), indicating that FACS intensity captured chromatin-state biology rather than cell-type composition. Heatmap clustering and Venn analysis likewise identified distinct transcriptional states across both HC and CFC conditions (**Fig. 5d,e**).

Gene-set analysis using a curated 240-gene hippocampal-plasticity set (**S. Table 1**) showed that CFC induced plasticity-associated gene transcription in both pools, however, the induction was stronger in H3K9me3^low^ neurons (Δ median log₂FC vs background +0.083, *P* < 0.0001) than in H3K9me3^high^ neurons (+0.066, *P* < 0.0001; interaction Δ median −0.04, *P* = 0.067). These data suggest that H3K9me3^high^ neurons engage plasticity programs less efficiently upon learning, which we next assessed at the pathway level. Pathway analysis of 38 curated pathway sets (MSigDB Hallmark, KEGG, GO BP, Reactome; **S. Table 2**) similarly showed stronger CFC-induced upregulation in H3K9me3^low^ neurons for 26 of 38 sets (**Fig. 5f**). Signed GSEA on the behavioral stimulation × H3K9me3-level interaction found reduced engagement of synaptic plasticity, glutamatergic synapse, long-term memory, calcium signaling, GABAergic synapse and long-term potentiation in H3K9me3^high^ neurons than in H3K9me3^low^ neurons (nominal *P* = 0.013–0.099, FDR > 0.25) (**Fig. 5g**). In contrast, dendritic-spine organization (*P* = 0.002, FDR = 0.062) and TGF-β signaling (*P* = 0.011, FDR = 0.096) were preferentially induced H3K9me3^high^ neurons consistent with synaptic stabilization rather than potentiation^31^. Together, these data suggest that H3K9me3^high^ neurons respond to learning by reinforcing existing synaptic architecture rather than mounting the activity-dependent signaling required for new plasticity.

Among 73 genes with significant condition x H3K9me3-level interactions, most changed in opposite directions after CFC in H3K9me3^high^ and H3K9me3^low^ neurons (**Fig. 5h, S**. **Table 3**). Nine (12%) were synaptic-plasticity genes, six of which were induced in H3K9me3^low^ neurons: Mapk1, Rps6ka2, Camk2d, Igsf11, Kdm6a, Itch. Together, these six form a memory-consolidation module spanning ERK/MAPK-CREB signaling (Mapk1, Rps6ka2), CaMKII signaling (Camk2d), AMPA-receptor stabilization (Igsf11), chromatin remodeling (Kdm6a), and ubiquitin-dependent protein turnover (Itch). In H3K9me3^high^ neurons, four of these genes were suppressed (Kdm6a, Mapk1, Igsf11, Rps6ka2) and Camk2d was unchanged, indicating a failure to engage this consolidation machinery. Importantly, while Itch was induced in both H3K9me3^high^ and H3K9me3^low^ neuronal pools, this gene was induced more strongly in H3K9me3^high^ neurons (Δ log₂FC +14.1 H3K9me3^high^ vs +7.6 H3K9me3^low^), demonstrating preserved transcriptional responsiveness and a selective, rather than floor, effect.

We next tested whether H3K9me3^high^ neurons were excluded from the engram, or instead engaged engram transcription without inducing plasticity machinery, using a canonical engram-UP core set (56 genes)^32^. Engram-UP programs were robustly induced by CFC in both populations. However, in H3K9me3^high^ neurons, this response was coupled to downregulation of kinase, AMPA-trafficking, and chromatin-opening machinery. In addition, GSEA of 38 transcription-factor target sets (TRRUST/ChEA; **S. Table 4**) showed that H3K9me3^high^ neurons preferentially engaged MEF2 target regulons (MEF2A/C/D, interaction FDR = 0.017–0.056), with Mef2a itself among the interaction hits. Because MEF2 transcription factors restrict synapse number^33^ during consolidation^34^, these findings indicate that H3K9me3^high^ neurons enter the engram while activating a program that constrains further plasticity.

Unsupervised analysis of all 1,386 significant transcripts supported a coordinated repression program. Weighted gene co-expression network analysis identified a 60-transcript module whose eigengene tracked the condition x H3K9me3 interaction (r = −0.91, *P* = 3.2 × 10⁻⁵). This module was selectively downregulated in CFC-H3K9me3^high^ neurons and unchanged in the other groups. The module included the interaction hits Adnp, Mta1 and Zfp972 alongside synaptic and signaling genes (Mef2c, Kalrn, Adam22, Cadm2, Rasgrp1; hubs Pik3r1, Acvr1c) (**Fig. S3b**).

Overall, the data reveal that dDG-H3K9me3^high^ neurons adopt a memory-stabilization transcriptional state that is locked in against plasticity-driven destabilization. Specifically, they engage the engram program and, rather than simply failing to induce the plasticity machinery, actively recruit synapse-restricting programs (MEF2 regulons and dendritic-spine/TGF-β stabilization) while suppressing the ERK/MAPK–CREB and chromatin-remodeling consolidation module. We next examined the functional consequences of dDG-H3K9me3^high^ neuron accumulation in aged mice.

### H3K9me3 accumulation impairs neuron activation and synaptic remodeling in aged dDG

Neuron activity and synaptic plasticity act as the two fundamental pillars of cognitive processes. Neuron activation is the substrate for engrams, the building blocks for memory^35^, while synaptic plasticity acts as the structural foundation, physically altering brain circuitry to encode and update memories ^36^. To directly test whether the age-related accumulation of H3K9me3 in dDG neurons alters both neuronal activation and protein content in synapses, we used our novel method NucleuSynapse-Tag^37^, which couples cell fractionation and flow cytometry to track neuronal activation, epigenetic status, and synaptic markers in parallel. To reduce H3K9me3 levels, Suv39h1 was knocked down by delivering AAV-EGFP-Suv39h1-shRNA viral particles in the dDG using stereotaxic surgery in aged mice (**Fig. S4a,b**). Suv39h1 knockdown was validated by quantifying Suv39h1 in both transfected (EGFP^+^) and non-transfected (EGFP^neg^) nuclei of dDG neurons (identified as NeuN^+^Prox1^+^) via flow cytometry. Compared to mice injected with control particles, mice injected with siRNA particles showed a reduced percentage of Suv39h1^+^ events (*P* = 0.007, **Fig. S4c,d**). For neuron activation, aged mice were briefly placed in a novel environment (NE), a naturalistic stimulus that activates a set of neurons and rapidly induces the expression of the immediate early gene Arc, which can be used as an activation marker^24^. As previous studies reported increased hippocampal protein synthesis following 3–4 h of NE-based paradigms^38,39^, animals were briefly placed in a NE (15 min) and returned to their home cage for 4 h (**Fig. 6a**), and immediately, nuclear and synaptosome fractions were obtained from dorsal hippocampi for NucleuSynapse-Tag analysis. A key methodological advantage of NucleuSynapse-Tag relies on the specific tracking of EGFP^+^ nuclei and synaptosomes derived from AAV-transfected neurons, via flow cytometry^37^. In mice injected with Suv39h1-siRNA particles, EGFP^+^ nuclei showed lower levels of H3K9me3 compared to EGFP^neg^ nuclei, in dDG neurons identified as NeuN^+^Prox1^+^ (*P* < 0.0001; **Fig. 6b,c**). Suv39h1 knockdown reduced the proportion of H3K9me3^high^ neurons (14.5±4.1% vs 36.7±6.5%, in EGFP^+^ and EGFP^neg^, respectively, *P* = 0.02; **Fig. 6b,d**), while it increased the proportion of the H3K9me3^neg^ population (35.1±6.2% vs 3.0±0.8%, in EGFP^+^ and EGFP^neg^, respectively, *P* = 0.0008; **Fig. 6b,d**). Notably, data showed a higher percentage of Arc^+^ events in EGFP^+^ than in EGFP^neg^ neurons in mice injected with Suv39h1-siRNA particles (*P* = 0.001, **Fig. 6e,f**). In contrast, mice injected with control particles showed comparable values of H3K9me3 (**Fig. 6b,c**), H3K9me3^neg^, H3K9me3^low^, and H3K9me3^high^ (**Fig. 6b,d**), and Arc^+^ nuclei in EGFP^neg^ and EGFP^+^ neurons (**Fig. 6e,f**). Overall, these results strongly support the idea that H3K9me3 accumulation reduces the capability of DG neurons to activate upon NE, thus unveiling a key functional impact of H3K9me3 in the aged brain.

**Fig. 6.**
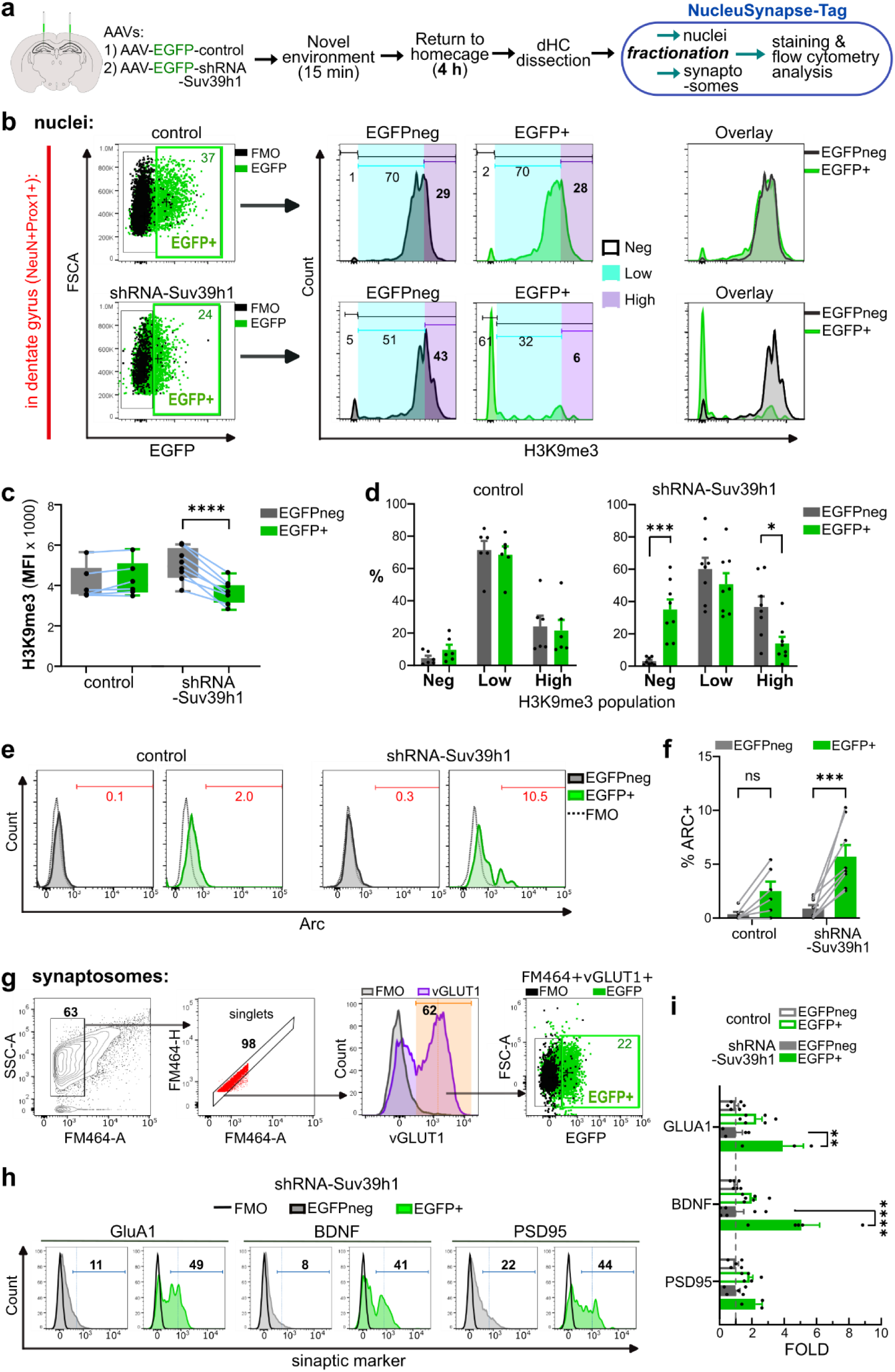
Suv39h1 RNAi reduces the proportion of H3K9me3^High^ neurons while increasing neuron activation and synaptic content of BDNF and GluA1 following NE in aged mice. **a,** AAV stereotaxic delivery in 18 m.o. mice was followed by placing animals in a novel environment (NE) for 15 min, then returning them to their cage for 4 h. The dorsal hippocampus was dissected to isolate, in parallel, nuclei and synaptosomes for immunostaining and flow cytometry analysis according to the NucleuSynapse-Tag protocol. **b,** Hierarchical gating strategy identified singlet DG nuclei by size and by co-staining for NeuN and Prox1 (not shown). Representative dot plots for gating on EGFP+ and EGFPneg events in DG (NeuN^+^Prox1+) nuclei (*left side*). Representative histograms (in the *center*) show the percentage of neurons H3K9me3^Low^ (blue area), H3K9me3^High^ (violet area), and H3K9me3^Neg^ (white area defined using an appropriate FMO staining control). **c,** Box plots show MFI (mean fluorescence intensity) of H3K9me3 samples of AAV-control (n = 6) and AAV-shRNA-Suv39h1 (n = 8) mice; *P* < 0.0001, Two-way ANOVA, main effect of EGFP-defined population (*P* < 0.0001), Sídák’s *post hoc* analysis: \*\*\*\**P* < 0.0001 (n = 14 mice). **d,** Bar graph shows % of H3K9me3-defined neuron subpopulations in AAV-control and AAV-shRNA-Suv39h1 mice; *P* < 0.0001, Two-way ANOVA, main effect of H3K9me3-defined population, Sídák’s *post hoc* analysis: \**P* = 0.02, \*\*\**P* < 0.0008. **e,** Representative histograms show the percentage of Arc^+^ events (red line) in EGFP^+^ and EGFP^neg^ events from DG (NeuN^+^Prox1^+^) nuclei. **f,** Bar graph showing % of Arc^+^ events in AAV-control and AAV-shRNA-Suv39h1 mice; *P* = 0.0001, Two-way ANOVA, main effect of EGFP-defined population, Sídák’s *post hoc* analysis: \*\*\**P* = 0.0001. **g,** Hierarchical gating for synaptosome analysis by flow cytometry (illustrated by arrows). Synaptosomes were stained with the lipophilic dye FM464 to trigger the detection of microparticles by flow cytometry. In the SSC-FM464 chart, the gating of synaptosomes focused on particles detected on the left side to discard large particles contaminated with aggregates. This was followed by doublet discrimination to select singlets, VGluT1^+^ events, and identifying synaptosomes from transfected neurons by EGFP. **h,** Representative histograms of protein levels of GluA1, BDNF, and PSD95 in synaptosomes from dorsal hippocampus of animals injected with shRNA-Suv39h1 AAV. **i,** Bar graph showing fold increase of percentage of positive events for each marker in EGFP+ relative to EGFPneg events in AAV-control (n = 6) and AAV-shRNA-Suv39h1 (n = 6) mice; \**P* < 0.0001, Two-way ANOVA main effect of AAV, Sidak’s *post hoc* analysis \*\**P* = 0.005, \*\*\*\**P* < 0.0001. Data are represented as mean ± s.e.m.

In parallel, NucleuSynapse-Tag allowed the quantification of synaptic markers in the synaptosome fraction of mice injected with viral particles. As we have previously reported^40-42^, we identified synaptosomes by size and FM4-64 (**Fig. 6g**), a membrane dye that binds to synaptosomes^37,43^. Next, we focused on excitatory synaptosomes by gating on vGLUT1^+^ events, which were then classified based on the EGFP signal as EGFP^neg^ and EGFP^+^ (**Fig. 6g**). Finally, we quantified the protein levels of GluA1, BDNF, and PSD95, proteins critical for plasticity and memory whose genes have been identified as H3K9me3 targets in the brain by genome-wide ChIP-seq and mRNA-seq^22,44,45^. Notably, synaptosomes from mice injected with Suv39h1-siRNA particles showed increased protein levels of GluA1 (*P* = 0.005) and BDNF (*P* < 0.0001) in EGFP^+^ relative to EGFP^neg^ synaptosomes, with no differences in PSD95 (**Fig. 6h,i**). In contrast, samples from mice injected with control particles showed comparable levels of BDNF, GluA1, and PSD95 in the EGFP^neg^ and EGFP^+^ populations (**Figs. 6h,i**). Thus, in the aged dDG, H3K9me3 accumulation impedes the increase of GluA1 and BDNF synaptic levels upon stimulation.

Next, we performed immunostaining on brain slices to further test the hypothesis that the age-related accumulation of H3K9me3^high^ dDG cells reduces the number of activated neurons after behavioral stimulation. H3K9me3 was genetically inhibited via Suv39h1-siRNA particles in the dDG of aged mice that were exposed to NE. Imaging of coronal brain slices was coupled to Image J-assisted analysis for single-cell quantification of Arc^+^ cells and H3K9me3 levels in EGFP^+^ and neighbor EGFP^neg^ cells in dDG (**Fig. 7a**). In mice injected with Suv39h1-siRNA particles, EGFP^+^ granule cells showed lower H3K9me3 levels (*P* = 0.0008, **Fig. 7b,c**) but a higher proportion of Arc^+^ events (*P* = 0.02, **Fig. 7d**), as compared to the neighbor EGFP^neg^ cells. In contrast, comparable levels of H3K9me3 and Arc^+^ neurons were found in EGFP^neg^ and EGFP^+^ populations in mice injected with control particles. Consistent with NucleuSynapse-Tag data (**Fig. 6e,f**), these findings support the idea that H3K9me3 accumulation reduces the pool of neurons available for activation in the dDG.

**Fig. 7.**
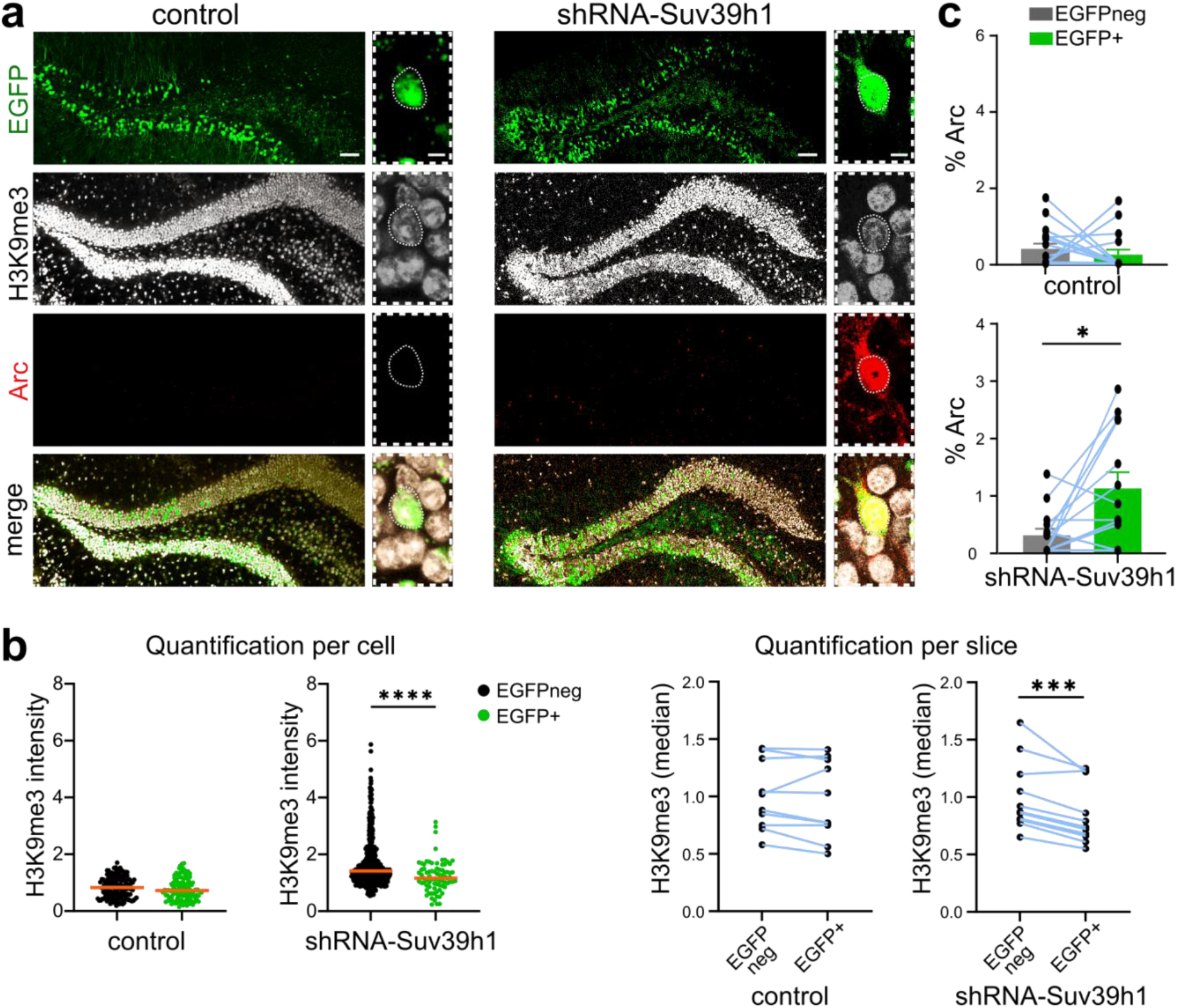
Genetic inhibition of Suv39h1 reduces H3K9me3 levels and increases Arc^+^ neurons in dorsal DG following NE in aged mice. AAV stereotaxic delivery in 18 m.o. mice was followed by placing the animals in a novel environment for 15 min, then returning them to their cage for 1 h. **a,** Coronal brain slices were immunostained for EGFP, H3K9me3, and Arc, scale bar = 50 μm; magnification, scale bar = 5μm. **b,** Intensity of H3K9me3 is shown per cell (n > 330/slice) for EGPF^+^ and neighbor EGFPneg cells for representative slices of control (n = 10 slices) and shRNA-Suv39h1 (n = 11 slices) groups. Median intensity for per-cell quantification is shown as a red line; *P < 0.0001, Wilcoxon test. Quantification per slice shows the median intensity of H3K9me3 in each slice of control (N = 4 mice) and shRNA-Suv39h1 (N = 4 mice) groups; *P = 0.0008, paired *t*-test. **c,** Percentage of Arc^+^ in EGFP^+^ and neighbor EGFPneg cells in control (N = 4 mice) and shRNA-Suv39h1 (N = 4 mice) groups. Paired dots show values in each slice, and underneath bar graphs show mean ± s.e.m., *P = 0.02, Wilcoxon test.

### Genetic inhibition of H3K9me3 in the dDG improves long-term memory in aged animals

To test whether genetic Suv39h1 inhibition suffices to restore cognitive function, a battery of memory tasks was performed in control- and siRNA-Suv39h1-injected mice (**Fig. 8a**), which showed comparable locomotor activity in the open field test, as evaluated by the total distance traveled inside an arena (**Fig. 8b,c**). Spatial long-term memory was evaluated in aged mice using the object location memory task (OLM) (**Figs. 8d**). Mice injected with Suv39h1-siRNA particles showed better performance compared to mice injected with control particles, as evaluated by the discrimination index (DI) (*P* = 0.03), whereas no significant differences were found in the training (acquisition) phase (**Fig. 8e,f**) or in total exploration times between groups (**Fig. S5a**). An independent set of aged mice was tested in the object recognition memory (ORM) task (**Fig. 8g**). Consistent with the OLM results, the group of animals injected with Suv39h1-siRNA particles showed a higher DI in the ORM test relative to control mice (*P* = 0.03), whereas both groups showed a similar DI in the acquisition phase (**Fig. 8h,i**) and similar total exploration times (**Fig. S5b**). A third group of aged mice was tested on the Y-maze task, another task used to assess spatial memory (**Fig. 8j**). Mice injected with siRNA particles showed better performance compared to control mice, as reflected by higher exploration times in the novel arm relative to the familiar arm (*P* = 0.001, **Fig. 8k**), with no differences found in total travelled distances or in total arm entries between groups (**Fig. S5c**), thus confirming an improvement in long-term memory by Suv39h1 knockdown. Two weeks later, animals were tested on contextual fear conditioning (CFC) (**Fig. 8l**). Both groups showed significant freezing behavior during the test relative to training (control *P* = 0.02; RNAi-Suv39h1 *P* = 0.003) (**Fig. 8m**). CFC induces a robust memory trace; therefore, it is not surprising that aged mice retained CFC memory, consistent with previous findings^9^. Overall, these data show improved long-term memory in aged mice when Suv39h1 is inhibited.

**Fig. 8.**
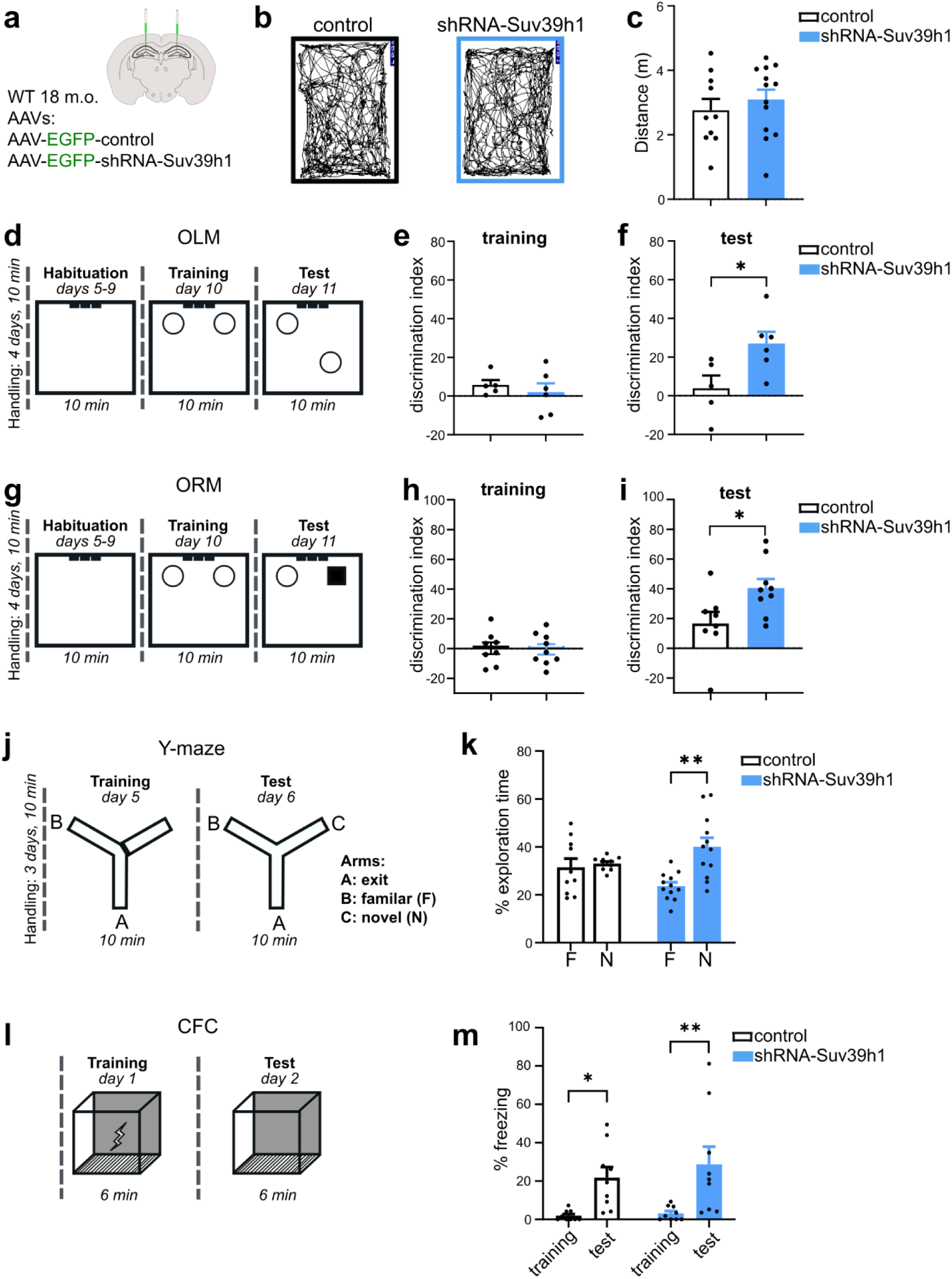
Suv39h1 knockdown in dDG improves long-term memory in aging male mice. **a,** 18-month-old mice with previous AAV stereotaxic delivery in the dorsal DG were used for all memory tasks. (**b,c**) Open field analysis shows representative trajectories (**b**) and total distance (**c**) traveled in a square arena (10-min test). (**d-f**) OLM task. **d,** Schematic showing the experimental design of the OLM task. **e,** No difference in the discrimination index (DI) during the OLM training session, measured by comparing time spent exploring both objects. **f,** DI in the test session (DI = ([time spent at novel object location — time spent at familiar object location]/[total time exploring both objects]) x100]); \**P* = 0.03, two-tailed unpaired *t*-test, n = 11 mice. (**g-i**) ORM task. **g,** Schematic showing the experimental design of the NOR task. **h,** DI during the ORM training session, measured by comparing time spent exploring both objects. **i,** DI comparing the novel object and the original object; \**P* =0.03, two-tailed unpaired *t*-test, n = 17 mice. (**j-l**) Y-maze task. **j,** Schematic showing the experimental design of the Y-maze task. **k,** Percentage of time spent in familiar and novel arms; *P* = 0.02, Two-way ANOVA, effect of arm-AAV, Sídák’s *post hoc* analysis: \*\**P* = 0.001, n = 22 mice. (**l,m**) CFC task. **l,** Schematic showing the experimental design of the CFC task. **m,** Freezing behavior in the training and test phases. *P* = 0.0003, Two-way ANOVA, main effect of test, Sídák’s *post hoc* analysis: \**P* = 0.02, \*\**P* = 0.004, n = 18 mice. Data are represented as mean ± s.e.m.

## Discussion

Epigenetic alterations are a hallmark of brain aging^46,47^. However, the impact of epigenetic mechanisms at the protein level and the corresponding functional consequences are poorly understood. Moreover, specific brain regions or cell types with high vulnerability to age-related epigenetic changes remain to be identified. Here, we used a highly quantitative approach at the single-event level with flow cytometry that allowed us to identify a subset of neurons that accumulates H3K9me3—via Suv39h1—in the hippocampus with age. A detailed analysis of hippocampal tissue demonstrates that H3K9me3^high^ neurons selectively emerge in the dDG. Nuclei sorting by FACS coupled to RNA-seq analysis shows that H3K9me3^high^ neurons from the dDG express a unique transcriptional signature linked to downregulation of plasticity-machinery genes. Using the NucleuSynapse-Tag protocol, we found that H3K9me3 accumulation interferes with both neuronal capacity for activation and synaptic remodeling in response to behavioral stimulation. Lastly, a decrease in the number of dDG-H3K9me3^high^ neurons by Suv39h1 knockdown results in improved long-term memory in aged mice, thus identifying Suv39h1 as a therapeutic target. Hence, based on molecular, cellular, and behavioral experiments, we identify a region-specific epigenetic mechanism that interferes with neuronal and synaptic activities in the aged hippocampus, thus impairing memory.

A growing body of evidence indicates that past experiences produce lasting changes in neurons via structured genomic responses, mainly controlled by epigenetic mechanisms^48^, including DNA methylation^49^ and histone posttranslational modifications^9,50^. Our findings suggest that the subset of dDG-H3K9me3^high^ neurons becomes less available for recruitment and unable to consolidate, supporting the idea that the epigenetic state determines neuronal eligibility for recruitment in the formation of a memory trace^51^ and retrieval^49^. In an elegant new study, it was shown that overexpression of histone acetyltransferases increased histone-3 acetylation at lysine 27 (H3K27AC) in amygdala neurons, leading to a primed chromatin state that was sufficient and necessary for memory allocation^51^. In the same study, it was further demonstrated that chromatin opening by H3K27AC improves memory by increasing neuronal excitability and synaptic density^51^. Similarly, our data provide evidence of an equivalent mechanism that works in the opposite direction, as we found that the age-dependent accumulation of H3K9me3, a mark associated with closed chromatin^29^, reduces neuron activation and the levels of plasticity-related proteins at the synapse, and impairs memory. To the best of our knowledge, no studies have evaluated how a *naturally occurring* epigenetic alteration modulates both neuron activation and synaptic protein remodeling, thereby linking molecular and functional consequences of H3K9me3 buildup in dDG neurons. Our results support a model in which intrinsic chromatin plasticity influences bidirectional and dynamic changes in neuronal activity, priming neurons for memory encoding. Although the full potential of H3K9me3 in regulating neuronal function, plasticity, and memory warrants further research, our study provides evidence of a repressive epigenetic mark negatively controlling memory processing in the dDG.

The hippocampus functions as a dynamic system that both isolates distinct experiences to prevent memory interference and stabilizes older, established memories^52^. This dual role is achieved through distinct mechanisms, including the generation of new neurons for encoding and the stabilization of synaptic changes^52^. Highlighting the role of epigenetics on hippocampal dynamics, recent evidence demonstrated that, in adult DG neurons, learning and neuronal activity trigger a late-onset increase in H3K9me3 levels that reduce wiring capacities but, surprisingly, contribute to the preservation of older memories^19^. Consistently, adult newborn DG neurons that are recruited in an engram undergo Suv39h1 and H3K9me3 upregulation, which reduces their ability to be re-recruited later, thus stabilizing stored memories^18^. Based on this evidence, we propose that the age-dependent accumulation of H3K9me3^high^ neurons—with diminished plasticity capabilities—would be the result of the process required to preserve information in stable circuits. Consequently, the pool of available neurons required to encode information in new circuitries would be cumulatively diminished with age, gradually changing the stability/plasticity equilibrium of neuronal circuits towards stability over plasticity. Our FACS-RNAseq data support this idea by showing that CFC induces a transcriptional state linked to reduced synaptic capability while reflecting memory stabilization specifically in H3K9me3^high^ neurons. Further, we found that reducing the amount of H3K9me3^high^ via Suv39h1 knockdown increases the number of Arc^+^ neurons in response to a novel environment and improves long-term memory in aged mice, defining SUV39H1 as a target for intervention.

The hippocampus is highly vulnerable to both normal aging and AD. Importantly, we found improvements in long-term memory tasks that strongly rely on intact hippocampal function (OLM and Y-maze) by genetic inhibition of H3K9me3 in the dDG. Hence, based on the discovery of dDG-H3K9me3^high^ neurons, our data provide evidence of a region-specific epigenetic mechanism that could impair the encoding of new memories in aged individuals, potentially contributing to anterograde amnesia, which occurs to a moderate-to-severe degree in the early stages of AD^53^. Although the contribution of H3K9me3^high^ neurons to anterograde amnesia in AD has not been formally tested, recent reports show that H3K9me3-mediated heterochromatin condensation is elevated in the cortex of mild cognitive impairment and sporadic AD^44^ postmortem brains with a strong negative correlation between H3K9me3 intensity and cognitive scores.

Genetic and pharmacological studies have consistently found that, via changes in chromatin accessibility at specific genomic regions and ultimately in gene expression^50^, histone posttranslational modifications act as critical starting points for regulating synaptic remodeling^18,19,44,54^. By exploring transcription patterns as a function of H3K9me3 neuronal levels in the dDG, our data show that H3K9me3^low^ and H3K9me3^high^ neurons express different genetic programs, with the H3K9me3^high^ population showing reduced learning-induced engagement of synaptic genes. Consistently, ChIP-seq and mRNA-seq analysis have previously identified synaptic genes, such as BDNF, as targets of H3K9me3 in the hippocampus of middle-aged^22^ and aged^20^ mice, and in the AD cortex^44^. Using NucleuSynapse-Tag coupled to Suv39h1 knockdown, our data indicate that excitatory (VGlut1^+^) synapses of dDG-H3K9me3^high^ neurons have reduced protein levels of BDNF and GluA1, which are crucial for synaptic plasticity^55^, including long-term potentiation^56^, a well-known cellular correlate of learning and memory^57,58^. Our data support the idea that epigenetic repression by H3K9me3 in dDG negatively impacts the transcription of synaptic genes and the content of plasticity-related proteins in synapses in response to behavioral stimulation in aged mice, thus linking cellular and molecular mechanisms on how epigenetic modifications regulate memory in aging.

Flow cytometry has become a valuable quantitative tool to study the complex cellularity of the central nervous system with high resolution, allowing high-content analysis at the single-event level of cells^59^, nuclei^24,37^, and synapses^37,40-42^. Using a flow cytometry-based approach, our cell-type-specific study reveals the presence of H3K9me3^high^ neurons, which became abundant in the dDG of aging mice. Our findings are supported by previous reports showing an age-dependent increase in H3K9me3 levels in olfactory neurons of *Drosophila*^60^, and in the retina^61^ and hippocampus of mice^20,21,62,63^, as assessed by semi-quantitative methods such as Western blot^21^ and immunohistochemistry^62,63^. Although our results indicate that Suv39h1 plays a fundamental role in H3K9me3 accumulation in dDG neurons with age, full characterization of the upstream mechanisms requires further study. H3K9 can be methylated by several HMTs, including G9a/GLA, which catalyzes H3K9 mono- and dimethylation; and by Suv39h1 and Suv39h2, which produce H3K9 trimethylation^29^. Conversely, H3K9me3 is demethylated by HDMs, such as the JMJD2 (KDM4) subfamily, which specifically targets H3K9me3^64,65^. Upstream of HMTs and HDMs, the activity-dependent phosphorylation of p66α by AMP-activated protein kinase recruits the histone deacetylase HDAC2 and Suv39h1 to produce H3K9me3, as shown in neuron cultures^19^. Therefore, based on previous reports, potential mechanisms for rising H3K9me3 levels in the mouse aged brain include an increase in the activity of AMP-activated protein kinase^66^ and in the expression of Suv39h1^21^, and a reduced expression of KDM3B (which demethylates H3K9me1/2) that leads to the accumulation of methylated H3K9 in genes for synapse assembly and vesicle transport, causing synaptic impairment^61^. In addition, a redistribution of methylated H3K9 from constitutive to facultative heterochromatin was observed in cells from aged retina, suggesting an age-related increase in repression of cell-type-specific genes besides the well-known physiological repression of developmental and non-tissue genes by H3K9me3 at constitutive heterochromatin^61^.

In summary, our findings indicate that what starts as a gradual epigenetic remodeling by H3K9me3 buildup in dDG neurons eventually contributes to neuron and synaptic dysfunction and, ultimately, to cognitive decline—but could potentially be treatable.

## Methods

Detailed descriptions of the experimental procedures and analytical methods are provided in the *Supplementary Methods*.

## Acknowledgments

The authors gratefully acknowledge the late Dr. Carl W. Cotman for his invaluable scientific advice, insightful guidance, and unwavering support throughout the development and execution of this research. We are grateful to Carina Uribe, Alejandra Castillo, and Christian Molina (Flow Cytometry Unit, LIIGH/UNAM) for technical assistance in flow cytometry experiments; María A. Carbajo and Alejandra Castilla (Vivarium/INB/UNAM) for laboratory animal management; and Jessica González Norris for manuscript editing. Larry Overman kindly provided ETP69. C.A.S.-R. is a student from the Programa de Maestría-Neurobiología (UNAM) and was supported by fellowships from CONAHCyT-México (CVU:1223128) and UNAM-DGAPA-PAPIIT IA203421 (#B221212). This work was supported by the US National Institute on Aging (R01 AG051807 and RF1 AG057558) to C.W.C., and by UNAM-DGAPA-PAPIIT (grants IA203421, IA208524 and IN226526), the Alzheimer’s Association (grant AARF-21-722869) and CONAHCyT (grant CF-2023-I-796) to G.A.P.

## Contributions

G.A.P. conceived and supervised the project. Y.M.L., C.J.R.O., and G.A.P. performed flow cytometry experiments. Y.M.L. performed IHC, stereotaxic injections, and NE, open field, ORM, Y-maze, and CFC tasks. C.J.R.O. performed FACS, RNA purification, stereotaxic injections, and the OLM task. Y.M.L. and G.A.P. performed NucleuSynapse-Tag experiments. C.S.R. performed NE and IHC. A.F.I. conducted all behavioral analyses. C.W.B. performed ETP injections and collected hippocampal tissue. N.C.B. performed RNA-seq bioinformatic analyses. G.A.P., C.J.R.O., and N.C.B. contributed to manuscript writing. All authors approved the final version.

## Ethics declarations

The authors declare no competing interests

## References

1. Lubin, F.D. Epigenetic gene regulation in the adult mammalian brain: multiple roles in memory formation. Neurobiology of learning and memory 96, 68–78 (2011).

2. Lattal, K.M. & Wood, M.A. Epigenetics and persistent memory: implications for reconsolidation and silent extinction beyond the zero. Nature neuroscience 16, 124–129 (2013).

3. Jarome, T.J. & Lubin, F.D. Epigenetic mechanisms of memory formation and reconsolidation. Neurobiology of learning and memory 115, 116–127 (2014).

4. Gräff, J., Woldemichael, B.T., Berchtold, D., Dewarrat, G. & Mansuy, I.M. Dynamic histone marks in the hippocampus and cortex facilitate memory consolidation. Nat Commun 3, 991 (2012).

5. Lardenoije, R., et al. The epigenetics of aging and neurodegeneration. Prog Neurobiol 131, 21–64 (2015).

6. Castellano, J.F., et al. Age-related memory impairment is associated with disrupted multivariate epigenetic coordination in the hippocampus. PLoS ONE 7, e33249 (2012).

7. Benevento, M., van de Molengraft, M., van Westen, R., van Bokhoven, H. & Kasri, N.N. The role of chromatin repressive marks in cognition and disease: A focus on the repressive complex GLP/G9a. Neurobiology of learning and memory 124, 88–96 (2015).

8. Gräff, J., et al. An epigenetic blockade of cognitive functions in the neurodegenerating brain. Nature 483, 222–226 (2012).

9. Peleg, S., et al. Altered histone acetylation is associated with age-dependent memory impairment in mice. Science 328, 753–756 (2010).

10. Disouky, A., et al. Human hippocampal neurogenesis in adulthood, ageing and Alzheimer’s disease. Nature 652, 1264–1273 (2026).

11. Shi, Y., et al. Histone demethylation mediated by the nuclear amine oxidase homolog LSD1. Cell 119, 941–953 (2004).

12. Parkel, S., Lopez-Atalaya, J.P. & Barco, A. Histone H3 lysine methylation in cognition and intellectual disability disorders. Learning & memory (Cold Spring Harbor, N.Y 20, 570–579 (2013).

13. Stewart, M.D., Li, J. & Wong, J. Relationship between histone H3 lysine 9 methylation, transcription repression, and heterochromatin protein 1 recruitment. Mol Cell Biol 25, 2525–2538 (2005).

14. Chaudhury, S., et al. FGF2 is a target and a trigger of epigenetic mechanisms associated with differences in emotionality: partnership with H3K9me3. Proceedings of the National Academy of Sciences of the United States of America 111, 11834–11839 (2014).

15. Hunter, R.G. Epigenetic effects of stress and corticosteroids in the brain. Front Cell Neurosci 6, 18 (2012).

16. Hunter, R.G., et al. Acute stress and hippocampal histone H3 lysine 9 trimethylation, a retrotransposon silencing response. Proc Natl Acad Sci U S A 109, 17657–17662 (2012).

17. Hunter, R.G., McCarthy, K.J., Milne, T.A., Pfaff, D.W. & McEwen, B.S. Regulation of hippocampal H3 histone methylation by acute and chronic stress. Proc Natl Acad Sci U S A 106, 20912–20917 (2009).

18. Chen, Q.N., Ding, X.L., Guo, X.X., Zhou, G. & Guan, J.S. Suv39h1 regulates memory stability by inhibiting the expression of Shank1 in hippocampal newborn neurons. Eur J Neurosci 55, 1424–1441 (2022).

19. Ding, X., et al. Activity-induced histone modifications govern Neurexin-1 mRNA splicing and memory preservation. Nat Neurosci 20, 690–699 (2017).

20. Snigdha, S., et al. H3K9me3 Inhibition Improves Memory, Promotes Spine Formation, and Increases BDNF Levels in the Aged Hippocampus. J Neurosci 36, 3611–3622 (2016).

21. Kushwaha, A. & Thakur, M.K. Increase in hippocampal histone H3K9me3 is negatively correlated with memory in old male mice. Biogerontology 21, 175–189 (2020).

22. Wu, T., et al. Histone H3K9 Trimethylation Downregulates the Expression of Brain-Derived Neurotrophic Factor in the Dorsal Hippocampus and Impairs Memory Formation During Anaesthesia and Surgery. Front Mol Neurosci 12, 246 (2019).

23. Fuchs, D.T., et al. H3K9me3 inhibition reverses Alzheimer’s progression by restoring synaptic and immune proteostasis across the brain-retina axis. Alzheimers Dement 22, e71918 (2026).

24. Jaeger, B.N., et al. A novel environment-evoked transcriptional signature predicts reactivity in single dentate granule neurons. Nature communications 9, 3084 (2018).

25. Bach, M.E., et al. Age-related defects in spatial memory are correlated with defects in the late phase of hippocampal long-term potentiation in vitro and are attenuated by drugs that enhance the cAMP signaling pathway. Proc Natl Acad Sci U S A 96, 5280–5285 (1999).

26. Dohm-Hansen, S., et al. The ‘middle-aging’ brain. Trends Neurosci 47, 259–272 (2024).

27. Boldrini, M., et al. Human Hippocampal Neurogenesis Persists throughout Aging. Cell Stem Cell 22, 589–599.e585 (2018).

28. Rapp, P.R. & Gallagher, M. Preserved neuron number in the hippocampus of aged rats with spatial learning deficits. Proc Natl Acad Sci U S A 93, 9926–9930 (1996).

29. Padeken, J., Methot, S.P. & Gasser, S.M. Establishment of H3K9-methylated heterochromatin and its functions in tissue differentiation and maintenance. Nat Rev Mol Cell Biol 23, 623–640 (2022).

30. Fanselow, M.S. & Dong, H.W. Are the dorsal and ventral hippocampus functionally distinct structures? Neuron 65, 7–19 (2010).

31. Mikheeva, I.B., Malkov, A.E., Pavlik, L.L., Arkhipov, V.I. & Levin, S.G. Effect of TGF-beta1 on long-term synaptic plasticity and distribution of AMPA receptors in the CA1 field of the hippocampus. Neurosci Lett 704, 95–99 (2019).

32. Rao-Ruiz, P., et al. Engram-specific transcriptome profiling of contextual memory consolidation. Nat Commun 10, 2232 (2019).

33. Flavell, S.W., et al. Activity-dependent regulation of MEF2 transcription factors suppresses excitatory synapse number. Science 311, 1008–1012 (2006).

34. Vetere, G., et al. Spine growth in the anterior cingulate cortex is necessary for the consolidation of contextual fear memory. Proc Natl Acad Sci U S A 108, 8456–8460 (2011).

35. Tonegawa, S., Morrissey, M.D. & Kitamura, T. The role of engram cells in the systems consolidation of memory. Nat Rev Neurosci 19, 485–498 (2018).

36. Takeuchi, T., Duszkiewicz, A.J. & Morris, R.G. The synaptic plasticity and memory hypothesis: encoding, storage and persistence. Philos Trans R Soc Lond B Biol Sci 369, 20130288 (2014).

37. Moreno-López, Y., Suárez-Rangel, C.A., Bonilla, J.A. & Prieto, G.A. Unveiling epigenetic-driven synaptic remodeling by parallel analysis of nuclei and synapses using NucleuSynapse-Tag. J Neurosci Methods 423, 110550 (2025).

38. Evans, H.T., Bodea, L.G. & Götz, J. Cell-specific non-canonical amino acid labelling identifies changes in the de novo proteome during memory formation. Elife 9(2020).

39. Monopoli, M.P., et al. Temporal proteomic profile of memory consolidation in the rat hippocampal dentate gyrus. Proteomics 11, 4189–4201 (2011).

40. Prieto, G.A., et al. Pharmacological Rescue of Long-Term Potentiation in Alzheimer Diseased Synapses. J Neurosci 37, 1197–1212 (2017).

41. Prieto, G.A., et al. Synapse-specific IL-1 receptor subunit reconfiguration augments vulnerability to IL-1β in the aged hippocampus. Proc Natl Acad Sci U S A 112, E5078–5087 (2015).

42. Prieto, G.A., Tong, L., Smith, E.D. & Cotman, C.W. TNFα and IL-1β but not IL-18 Suppresses Hippocampal Long-Term Potentiation Directly at the Synapse. Neurochem Res 44, 49–60 (2019).

43. Daniel, J.A., Malladi, C.S., Kettle, E., McCluskey, A. & Robinson, P.J. Analysis of synaptic vesicle endocytosis in synaptosomes by high-content screening. Nat Protoc 7, 1439–1455 (2012).

44. Lee, M.Y., et al. Epigenome signatures landscaped by histone H3K9me3 are associated with the synaptic dysfunction in Alzheimer’s disease. Aging Cell 19, e13153 (2020).

45. Bustos, F.J., et al. Epigenetic editing of the Dlg4/PSD95 gene improves cognition in aged and Alzheimer’s disease mice. Brain 140, 3252–3268 (2017).

46. Oberdoerffer, P., et al. SIRT1 redistribution on chromatin promotes genomic stability but alters gene expression during aging. Cell 135, 907–918 (2008).

47. Pal, S. & Tyler, J.K. Epigenetics and aging. Sci Adv 2, e1600584 (2016).

48. Mews, P., et al. From Circuits to Chromatin: The Emerging Role of Epigenetics in Mental Health. J Neurosci 41, 873–882 (2021).

49. Gulmez Karaca, K., et al. Neuronal ensemble-specific DNA methylation strengthens engram stability. Nat Commun 11, 639 (2020).

50. Mews, P., et al. Acetyl-CoA synthetase regulates histone acetylation and hippocampal memory. Nature 546, 381–386 (2017).

51. Santoni, G., et al. Chromatin plasticity predetermines neuronal eligibility for memory trace formation. Science 385, eadg9982 (2024).

52. Miller, S.M. & Sahay, A. Functions of adult-born neurons in hippocampal memory interference and indexing. Nat Neurosci 22, 1565–1575 (2019).

53. Jack, C.R., et al. Revised criteria for diagnosis and staging of Alzheimer’s disease: Alzheimer’s Association Workgroup. Alzheimers Dement 20, 5143–5169 (2024).

54. Fujiwara, K., et al. Deletion of JMJD2B in neurons leads to defective spine maturation, hyperactive behavior and memory deficits in mouse. Transl Psychiatry 6, e766 (2016).

55. Chen, L.Y., Rex, C.S., Pham, D.T., Lynch, G. & Gall, C.M. BDNF signaling during learning is regionally differentiated within hippocampus. J Neurosci 30, 15097–15101 (2010).

56. Muller, D., Joly, M. & Lynch, G. Contributions of quisqualate and NMDA receptors to the induction and expression of LTP. Science 242, 1694–1697 (1988).

57. Fedulov, V., et al. Evidence that long-term potentiation occurs within individual hippocampal synapses during learning. J Neurosci 27, 8031–8039 (2007).

58. Whitlock, J.R., Heynen, A.J., Shuler, M.G. & Bear, M.F. Learning induces long-term potentiation in the hippocampus. Science 313, 1093–1097 (2006).

59. Lobo, M.K., Karsten, S.L., Gray, M., Geschwind, D.H. & Yang, X.W. FACS-array profiling of striatal projection neuron subtypes in juvenile and adult mouse brains. Nat Neurosci 9, 443–452 (2006).

60. Muñoz-Carvajal, F., Sanhueza, N., Sanhueza, M. & Court, F.A. Age-dependent H3K9 trimethylation by dSetdb1 impairs mitochondrial UPR leading to degeneration of olfactory neurons and loss of olfactory function in. Elife 15(2026).

61. An, M.J., et al. Reorganization of H3K9me heterochromatin leads to neuronal impairment via the cascading destruction of the KDM3B-centered epigenomic network. iScience 27, 110380 (2024).

62. Kim, H., et al. Bifidobacterium bifidum BGN4 and Bifidobacterium longum BORI promotes neuronal rejuvenation in aged mice. Biochem Biophys Res Commun 603, 41–48 (2022).

63. Wu, Z.X., et al. Accelerated Deficits of Spatial Learning and Memory Resulting From Prenatal Inflammatory Insult Are Correlated With Abnormal Phosphorylation and Methylation of Histone 3 in CD-1 Mice. Front Aging Neurosci 11, 114 (2019).

64. Chen, Z., et al. Structural insights into histone demethylation by JMJD2 family members. Cell 125, 691–702 (2006).

65. Whetstine, J.R., et al. Reversal of histone lysine trimethylation by the JMJD2 family of histone demethylases. Cell 125, 467–481 (2006).

66. Wang, B.Z., Yang, J.J., Zhang, H., Smith, C.A. & Jin, K. AMPK Signaling Regulates the Age-Related Decline of Hippocampal Neurogenesis. Aging Dis 10, 1058–1074 (2019).

