## Supplemental Figures for "An H3K9me3^high^ epigenetic state constrains neuronal activation and synaptic remodeling in the aged dentate gyrus"

**Supplementary figures**

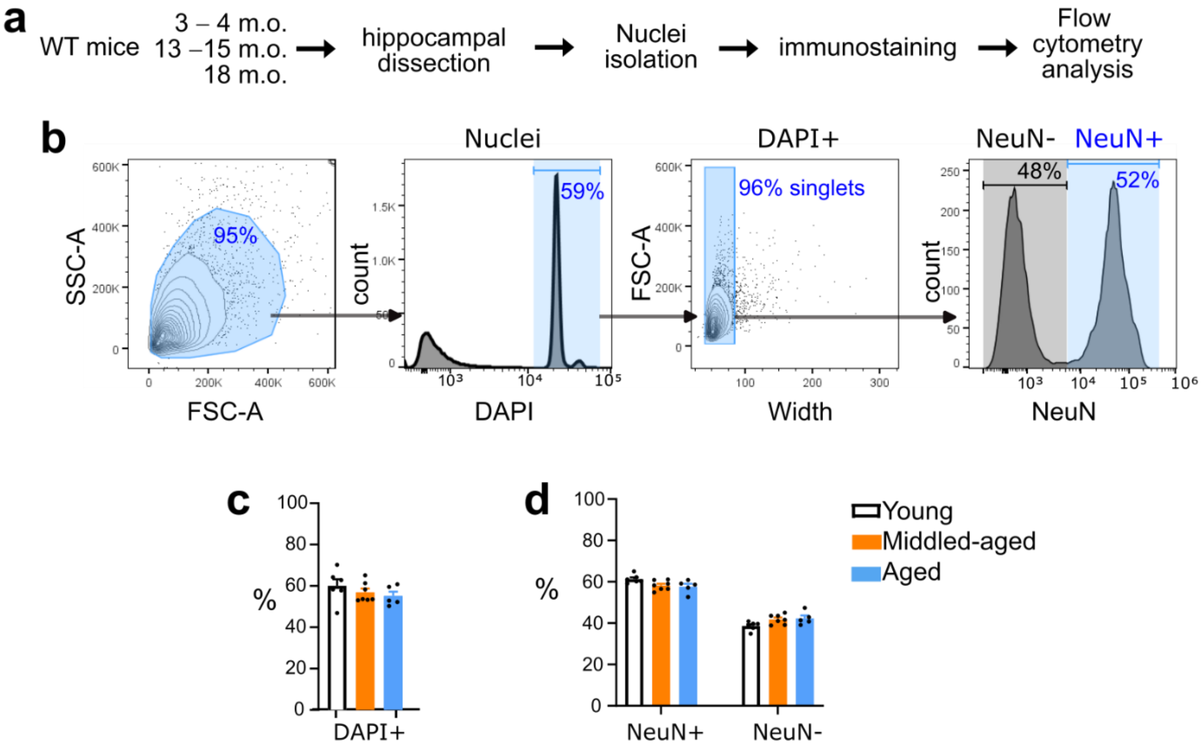


**Fig. S1. Comparable number of DAPI$^{+}$, NeuN$^{+}$ and NeuN$^{neg}$ events across age in the mouse hippocampus.** **a,** Workflow: WT mice at 3-, 15-, and 18-month-old (young, middle-aged, and aged, respectively) were used for flow cytometry analysis of cell nuclei from hippocampi. **b,** Representative FACS plots show the hierarchical gating strategy to identify nuclei by size (FSC-SSC plot) and DAPI staining and further distinguish glia and neurons by NeuN labeling in singlets (events selected by doublet discrimination). **c,** Bar graph shows similar percentages of DAPI$^{+}$ events in hippocampal neurons of young (n = 6), middle-aged (n = 7) and aged (n = 5) mice; $P$ = 0.403, ANOVA. **d,** Bar graph shows the percentage of NeuN$^{+}$ and NeuN$^{—}$ events; $P$ = 0.999, effect of "age", Two-way ANOVA. Data are represented as mean ± s.e.m.

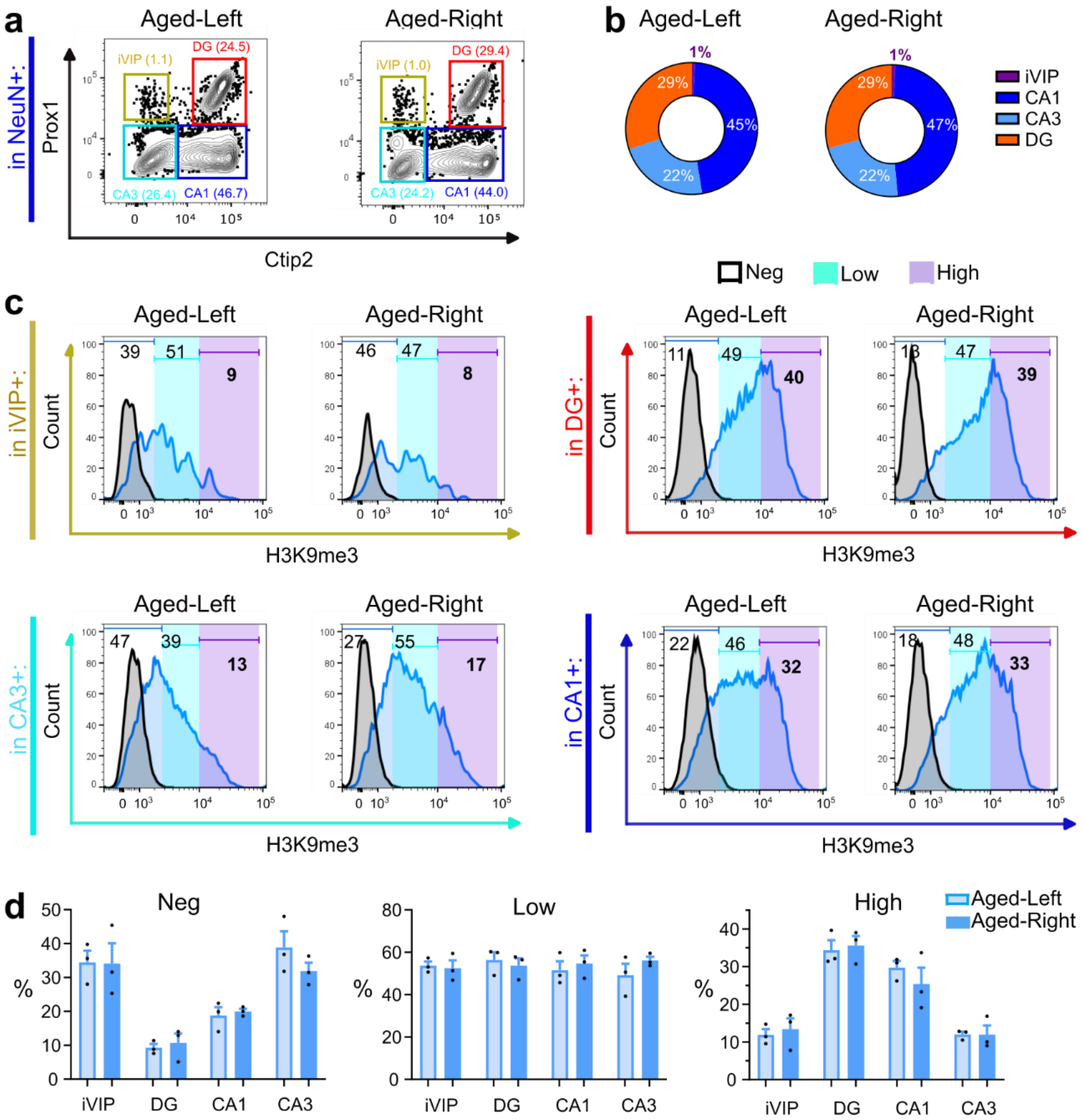


**Fig. S2. Comparable levels of H3K9me3$^{High}$ neurons in right and left hippocampi of aged mice. a,** Flow cytometry dissection of the hippocampus using Ctip2 and Prox1 markers in NeuN$^{+}$ nuclei obtained from dissected dorsal and ventral regions. Representative FACS plots show iVIP, DG, CA1 and CA3 populations (% are shown in parentheses). **b,** Circle charts show comparable proportions (%) of iVIP, DG, CA1, and CA3 populations in right and left hippocampi. **c,** Representative histograms of H3K9me3 levels in iVIP, DG, CA1, and CA3 neurons in right and left sides. **d,** Percentage of H3K9me3$^{Neg}$, H3K9me3$^{Low}$ and H3K9me3$^{High}$ neurons in iVIP, DG, CA1, and CA3 subfields, comparing right and left hippocampi in aged (n = 3) mice. No differences were observed: $P = 0.619$ (H3K9me3$^{Neg}$), $P = 0.282$ (H3K9me3 $^{Low}$), $P = 0.767$ (H3K9me3 $^{High}$), Two-way ANOVA. Data are represented as mean ± s.e.m.

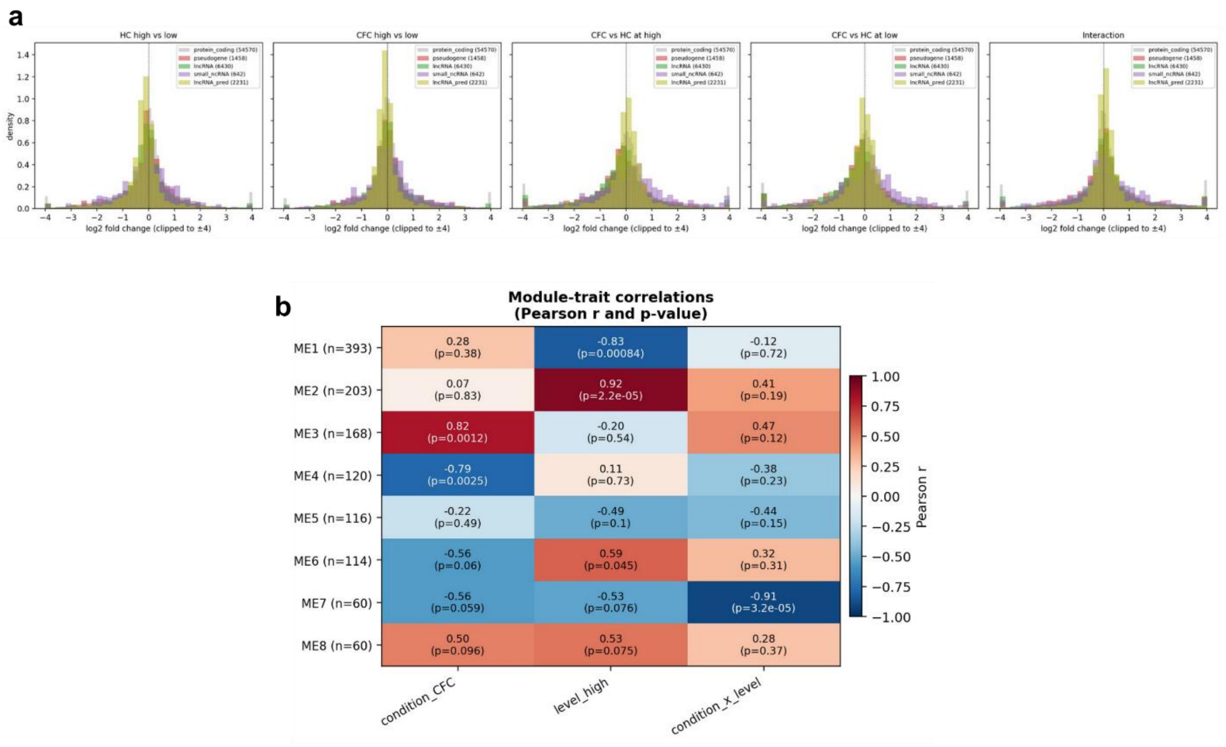


**Fig. S3. Biotype-stratified fold-change distributions and co-expression module–trait correlations. a,** $Log_2$ fold-change (LFC) distributions by transcript biotype for each of the five contrasts of the nested paired model. Grey histogram, protein-coding background (n = 54,570 transcripts); coloured overlays, pseudogenes (n = 1,458), lncRNAs (n = 6,430), small ncRNAs (n = 642) and predicted lncRNAs (n = 2,231); LFCs are clipped at ±4 for display. In the homecage high-versus-low contrast, the non-coding classes canonically silenced by H3K9me3 are shifted to lower expression in the H3K9me3$^{high}$ pool relative to the protein-coding background (pseudogenes, median LFC −0.053 versus +0.023; lncRNAs, −0.072; Mann–Whitney $P < 10^{-10}$), providing a data-internal positive control that FACS-defined H3K9me3 intensity reflects functional H3K9me3-mediated repression rather than cell-type composition. **b,** WGCNA module–trait correlation matrix. Unsupervised co-expression modules (ME1–ME8; module size in parentheses, transcripts) were correlated with binary sample traits: condition (CFC = 1), H3K9me3 level (high = 1) and their interaction. Cells give Pearson r (top) and P (bottom; n = 12 samples), coloured by r. ME2 tracks H3K9me3 level ($r = 0.92$, $P = 2.2 \times 10^{-5}$) and ME3/ME4 track condition ($r = 0.82$ and $-0.79$), whereas ME7 (n = 60 transcripts) is the only module tracking the condition × level interaction ($r = -0.91$, $P = 3.2 \times 10^{-5}$), consistent with selective downregulation of this module in CFC-H3K9me3$^{high}$ neurons.

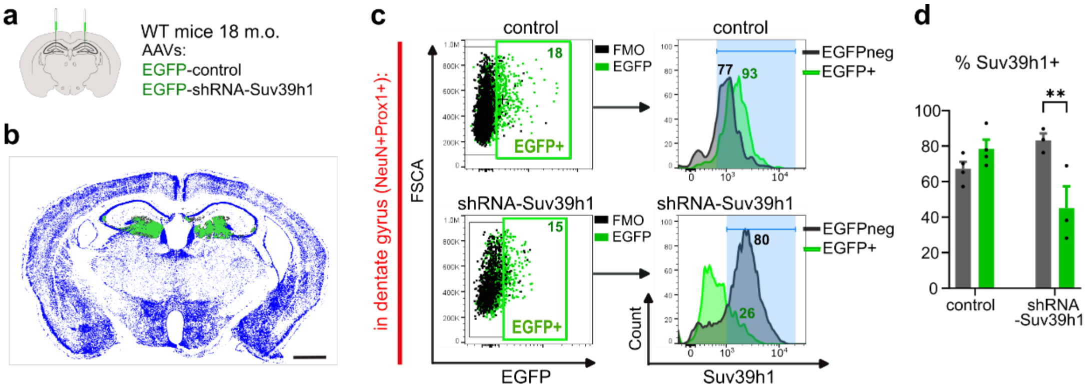


**Fig. S4. Suv39h1 knockdown validation. a,** After two months of AAV stereotaxic surgery in 18-month-old (m.o.) mice, site injection and Suv39h1 knockdown were verified. **b,** Coronal brain slice showing EGFP immunostaining; scale bar = 1000 µm. **c,** Representative dot plots (*left side*) for gating on EGFP$^{+}$ and EGFPneg events in DG (NeuN$^{+}$Prox1$^{+}$) nuclei, and histograms (*right side*) showing the percentage of Suv39h1$^{+}$ events; the blue area was set based on an appropriate FMO staining control. **d,** Bar graph shows % of Suv39h1$^{+}$ events in AAV-control (n = 4) and AAV-shRNA-Suv39h1 (n = 3) mice; $P$ = 0.004, Two-way ANOVA, Sídák's *post hoc* analysis: **$P$ = 0.007.

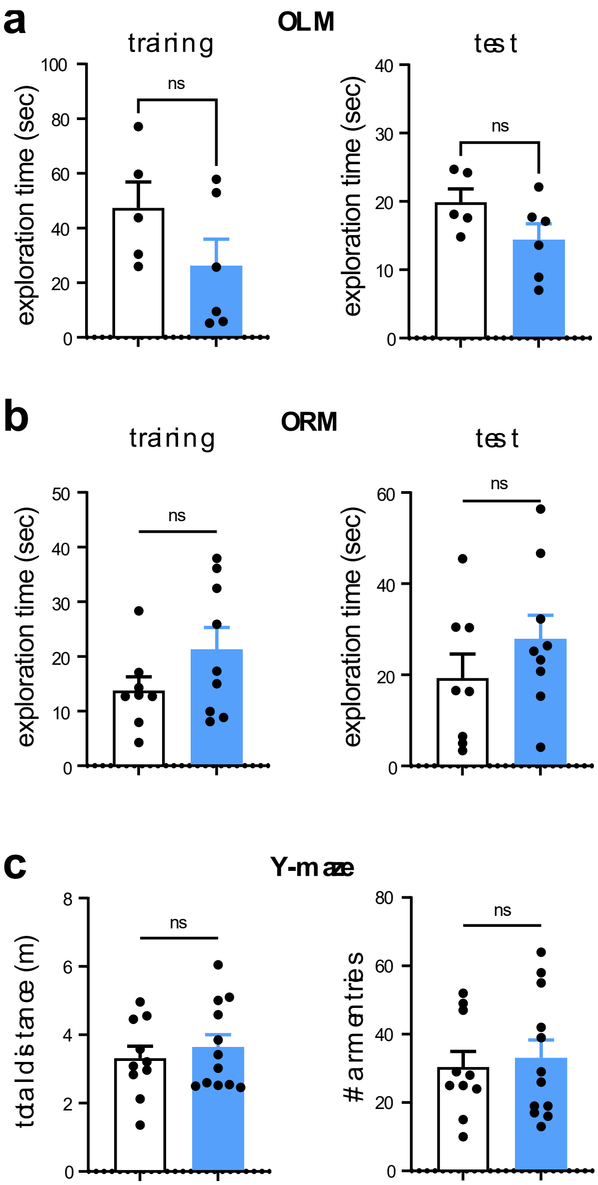


**Fig. S5. Total exploration times, distances, and arm entries in memory tasks following Suv39h1 knockdown in aging male mice.** We used 18-month-old mice with previous AAV stereotaxic delivery in the dorsal DG for all memory tasks. **a,** Total exploration times in training and test in the OLM task; n = 11 mice. **b,** Total exploration times in training and test in the ORM task; n = 17 mice. **c,** Total distances and total arm entries in the Y-maze task; n = 22 mice. Data are represented as mean ± s.e.m.; ns = not significant.
