## Supporting information (METHODS) for "An H3K9me3^high^ epigenetic state constrains neuronal activation and synaptic remodeling in the aged dentate gyrus"

### Animals

Male C57BL/6N mice were used at 3-, 15-, and 18-month-old. Mice were housed in groups of 2–3 per cage in acrylic cages with food and water ad libitum, at 30–70% humidity and 20–26°C. Lights were maintained on a 12:12 light/dark cycle. All procedures in this study followed the Principles of Laboratory Animal Care from the NIH and were approved by the bioethics committee at the Instituto de Neurobiología-UNAM (Protocol 121.A) and by the University of California, Irvine, Institutional Animal Care and Use Committee.

### Cell nuclei isolation

Acute hippocampal slices were obtained from mice (3-, 15-, and 18-month-old) as follows: mice were anesthetized with $CO_2$ (flow rate 30%) for 1 min, decapitated, and the brain quickly submerged in ice-cold PBS for 10 sec. Immediately, hippocampi were rapidly dissected on a cold surface; for some experiments, the hippocampus was split into dorsal and ventral regions. A McIlwain tissue chopper was used to slice (500-µm-thick) the hippocampus. Slices were transferred to a tube containing 1.2 mL of 1X Fix solution (True-Nuclear Transcription Factor Buffer Set, Biolegend, #424401), and incubated for 15 min in rotation, light-protected (RT). Fixation was quenched by removing Fix solution by vacuum (slices sink by gravity) and by adding 1 mL of the nuclear isolation buffer (NIB) (in mM: sucrose, 50; KCl, 25; MgCl2, 5; NH4Cl, 50; and Tris, pH 7.4, 120) (PMID: 24713002), which contains the formaldehyde-neutralizing chemical $NH4^+$ (in mM: sucrose, 50; KCl, 25; MgCl2, 5; NH4Cl, 50; and Tris, pH 7.4, 120). Slices were washed twice with 1.5 mL of NIB. Cross-linked tissue was homogenized in a 2-mL Dounce homogenizer (Sigma, D8938) containing 1 mL of NIB. Twenty-five strokes of pestle A (clearance 76–127 µm) followed by 25 strokes of pestle B (clearance 13–63 µm) were applied gently to avoid damage to nuclei (PMID: 24713002). After homogenization, large debris was removed by filtering hippocampal extracts using pre-wet MACS® SmartStrainers (30 µm) (Miltenyi, #130–098–458).

### Immunostaining of isolated nuclei

The protein concentration in the enriched nuclear fraction was determined by the BCA assay (ThermoFisher) using BSA standards. A volume containing 100 µg of nuclear fraction was transferred into cytometry tubes, and 2 mL of the Perm/Wash Buffer 1X (Biolegend, #424401) was added to each tube. All tubes were centrifuged at 2000 × g for 5 min at 20°C using a swinging rotor (5910R centrifuge, Eppendorf). After centrifugation, the supernatant was discarded by total inversion of the tube, and any excess liquid was gently dried with a paper towel while the tube remained inverted. Next, the pellet was resuspended through gentle finger agitation (no vortexing) before adding 250 µl of Perm/Wash Buffer 1X with primary antibodies. Tubes were incubated for 1 h (RT) in shaking. Primary antibodies were washed with Perm/Wash Buffer 1X followed by centrifugation (2000 × g, 20°C) for 5 min. Supernatant was discarded, and the pellet was incubated with a mix of secondary antibodies in 250 µl of Perm/Wash Buffer 1X for 30 min, in shaking and in the dark (RT). Samples were washed with 0.1 % Triton-X and centrifuged (2000 × g, 20°C) for 5 min. The supernatant was discarded, and the pellet was resuspended in 350 μL of PBS for flow cytometry analysis.

### Flow cytometry

Data were acquired using a NovoCyte 3000 cytometer (ACEA) equipped with violet (405), blue (488nm), and red (640nm) lasers. Fluorophores and dyes were detected by coupling lasers with specific channels (nm): 445/45, 530/30, 585/40, 675/30. Cross-reactivity was prevented by using primary antibodies from different hosts and/or different isotypes (*see Antibodies panels*), and by using isotype-specific (highly cross-adsorbed) secondary antibodies. Instrument settings, gating strategies, and controls for background fluorescence followed standard protocols for flow cytometry (Baumgarth and Roederer, 2000; Hulspas et al., 2009). Cytometer settings were as follows: threshold in FSC-H =125,000; stop criteria = 100,000 events or 250μl; laser intensity adjusted based on fluorescence controls. For hierarchical gating, the nuclei-containing region was established based on initial experiments locating DAPI-positive events in the FSC-SSC plot. Doublets were excluded using FSC-A *versus* FSC-W or FSC-H *versus* FSC-A plots. Fluorescence Minus One (FMO) staining controls were used to define negative and positive signals; in particular, the FMO for EGFP was prepared using ventral hippocampus (with no AAV delivery). Fluorescence spillover was corrected by compensation, using single-stained samples to generate compensation matrices. All antibodies were titrated. The cut-off value for defining the H3K9me3$^{high}$ subpopulation was rigorously set in the BL2 fluorescence channel (detecting phycoerythrin, PE). Data analysis was performed using FlowJo V.11 software (LLC).

### ETP injections

Mice received a single intraperitoneal injection of ETP69 (1 mg/kg) or vehicle (DMSO in saline 1:1) between 10:00 and 11:00 am. After 24 h, hippocampi were dissected and cryopreserved in controlled-rate freezing containers (CoolCell, Corning) at -72°C in sucrose buffer (320 mM sucrose, 10 mM HEPES, pH = 7 with NaOH). For cytometry analysis, samples were rapidly thawed (10-12 min in a water bath at 37°C) for nuclei isolation and immunostaining.

### Brain slice immunostaining and image acquisition

Mice were deeply anesthetized with phenobarbital (60–90 mg/kg) and transcardially perfused with saline solution (0.9% NaCl) followed by 4% PFA in saline. Brains were maintained in 4% PFA (4◦C) overnight and then cryoprotected in 30% sucrose in KPBS (potassium phosphate-buffered saline) for 48 h (4°C). Brains were frozen and sectioned (30 μm) using a CM1510 cryostat (LEICA). Brain slices were rinsed with KPBS, pretreated with 0.3% $H_2O_2$ and 0.1% $NaBH_4$, and permeabilized with 1% Triton X-100 for 30 min (RT). Primary antibody incubation (0.5% Triton X-100, 4°C, with shaking for 48 h) was followed by washing and secondary antibody incubation (0.5% Triton X-100, RT, with shaking for 2 h). Slices were mounted on a glass slide using VECTASHIELD antifade mounting medium. Primary antibodies used were rat anti-GFP Alexa

Fluor 488 (1:200, #338008, Biolegend), rabbit IgG anti-H3K9me3 (1:500, #ab8898, Abcam) and mouse IgG2a anti-Arc (sc-17839, Santa Cruz). Secondary antibodies: goat anti-rabbit IgG Alexa Fluor 647 (1:500, #A21245, Invitrogen) and goat anti-mouse IgG Alexa Fluor 555 (1:500, #A21422, Invitrogen). Confocal images were acquired on a Zeiss LSM 780 Duo confocal microscope. Whole brain sections were imaged using a 10× objective and the TileScan tool. Images of the dentate gyrus were acquired using a 40× objective and the TileScan tool. Linear unmixing using the ZEN software separated overlapping fluorescence signals. Single-cell image analysis was performed using Fiji (version 1.4f). The mean fluorescence intensity (MFI) of H3K9me3 was quantified in each nucleus, which was classified as $EGFP^+$ or $EGFP^-$, and $ARC^+$ or $ARC^-$ based on fluorescence intensities in appropriate detectors.

### Virus injection

Mice (18-month-old) were deeply anesthetized with isoflurane: 5 % for induction, 1–2 % for surgery. The animal's head was positioned in a stereotaxic instrument, secured with ear bars and stabilized with a nose restrainer. Eye lubricant was applied to both eyes to prevent drying, and 0.1 mL of 2% lidocaine was infiltrated into the incision site. The scalp was cleaned with iodine solution and 70 % ethanol for the craniotomy (1 mm). Coordinates for injection: AP =-1.6, ML =±0.9 mm, and depth =-2.2 mm. A bilateral injection of 150 nl AAV2-CAG-EGFP (Addgene # 37825, $1\times10^{13}$ vg/mL), AAV2-hU6-mSUV39H1-SCRAMBLE-shRNA-CAG-EGFP or AAV2-hU6-mSUV39H1-shRNA-CAG-EGFP (AAVnerGene Inc., $1 \times 10^{13}$ gc/mL) was performed using a pulled glass pipette connected to a thin plastic tube attached to a 20 mL syringe, as previously reported (Cetin et al., 2006). The pipette was kept in place for an additional 7 min to prevent backflow of the solution (Cetin et al., 2006). Mice received ibuprofen, 40 mg/300 mL in their water bottles for 5 days following surgery.

### Novel environment exposure

For novel environment (NE) exposure, all animals in a cage (2–3) were transferred to an acrylic cage (53 ×43 ×19 cm) with 2 cm of sawdust bedding and various objects: tunnels (2), Legos (3), balls (4), and a climbing wall, all in different colors. The enrichment items and cage were cleaned with Extran (Merck) and water before the experiment. The NE session lasted 15 min. For the one-hour readout experiments (Fig. 6), mice were placed in the NE between 9:00 and 9:30 a.m., returned to their home cages, and euthanized between 10:00 and 10:30 a.m. for hippocampal dissection and processing. For the four-hour readout experiments (Fig. 7), mice were placed in the NE between 7:00 and 7:30 a.m., returned to their home cages, and euthanized between 10:00 and 10:30 a.m., during the light cycle.

### Object location memory (OLM)

Mice were handled for four days (two sessions of 5 min/day) before starting habituation to the arena. Each mouse was habituated for five consecutive days to a white acrylic 5-sided box (40 cm × 30 cm × 30 cm) with soft illumination. Three black vertical stripes (25 cm, 1.9 cm wide, spaced by 1.5 cm) were fixed in the center of one of the walls of the arena, and the floor was covered with asser bedding. On training (day 10), mice were exposed to two identical objects (100-ml beakers filled with cement) placed at opposite ends of the arena for 10 min. On test (24 hr later, day 11), one of the objects was moved to a novel location within the arena, and mice were allowed to explore for 10 min. Objects and their relative positions were counterbalanced, and, to avoid olfactory cues, objects were thoroughly cleaned with 70% ethanol, and bedding was stirred after each trial. A video camera was mounted above the arena, and all sessions were recorded. Videos were hand-scored offline by a trained experimenter blind to the animal's IDs and the corresponding AAV treatment. The discrimination index was calculated using the formula: [(time exploring the object in the novel location– time exploring the object in the familiar location)/(total exploration time)] × 100. Exploration was considered as pointing the head toward an object. Turning around, chewing, or sitting on the objects was not considered exploratory behavior.

**Object recognition memory (ORM)**

Mice were handled for four days (two sessions of 5 min/day) before starting habituation to the arena. Each mouse was habituated for five consecutive days to a white acrylic 5-sided box (40 cm × 30 cm × 30 cm) with soft illumination. On training (day 10), mice were exposed to two identical objects (100-ml beakers filled with cement) placed at opposite ends of the arena for 10 min. On test (24 hr later, day 11), one of the objects was replaced with a novel object (a white light bulb), and mice were allowed to explore for 10 min. Objects were counterbalanced, and to avoid olfactory cues, objects were thoroughly cleaned with 70% ethanol, and bedding was stirred after each trial. A video camera was mounted above the arena, and all sessions were recorded. Videos were hand-scored offline by a trained experimenter blind to the animal's IDs and the corresponding AAV treatment. The discrimination index was calculated using the formula: [(time exploring the novel object – time exploring the familiar)/(total exploration time)] × 100. Exploration was considered as pointing the head toward an object. Turning around, chewing, or sitting on the objects was not considered exploratory behavior. Total distances covered by each animal were also quantified, as a measure of locomotor activity.

**Y-maze task**

Mice were handled for three days (two sessions of 5 min/day). Before the trial, mice were habituated to the testing room environment for 30 minutes. The apparatus, made of white-opaque acrylic, consisted of three identical arms (50 cm length × 8.5 cm width × 25 cm height) symmetrically diverging at 120° angles from a central, triangular platform, with soft illumination. The arms, designated A, B, and C, contained internal and external cues (posters and shapes on the walls). During training, mice were placed in the maze with one arm closed off (novel arm) and allowed to explore for 10 min. On test (24 hr later), mice were placed back in the start arm with all three arms fully accessible. A video camera was mounted above the maze, and all sessions were recorded. A computerized tracking system SMART 3.0 (Panlab) was used to quantify the percentage of time spent on each arm and to calculate the spontaneous alternation percentage, defined as the number of consecutive triplet entries (i.e., entering three different arms, such as A-B-C or C-B-A) divided by the total number of arm entries minus two, multiplied by 100. Total arm entries and total distances covered by each animal were also quantified, as a measure of locomotor activity.

#### Contextual fear conditioning (CFC)

CFC was conducted in a standard fear conditioning chamber (San Diego Instruments) equipped with a grid floor (Context A: black circles on white walls). The chamber was cleaned with 70% ethanol between subjects. During training, mice were placed in the chamber and allowed to explore for 3 min. Following this acclimation period, a mild footshock (0.2 mA, 1 sec) was delivered; a second shock (0.2 mA, 1 sec) was delivered after 3min. Animals remained in the chamber for an additional 3 min before being returned to their home cages. On test (24 hr later), mice were returned to the same conditioning chamber (Context A). A computerized tracking system SMART Anymaze was used to quantify the freezing behavior over the 6-minute test period and compared to the 3 min period before the first shock on the training day.

#### Open field test

Mice were placed in the periphery of a square arena (40 cm × 30 cm × 30 cm) with soft illumination and recorded for 10 min via an overhead video tracking system. SMART-based analysis was used for measuring total traveled distance.

#### NucleuSynapse-Tag

The NucleuSynapse-Tag protocol has been recently described in detail (PMID: 40835005). Briefly, mice were deeply anesthetized with $CO_2$ before brain extraction. The brain was placed in ice-blended PBS for 5–10 s. All materials, including solutions, grinder, pestle, and tubes, were precooled on ice for tissue dissection and homogenization. Dorsal hippocampi were rapidly dissected on an ice-cold surface and sectioned with a knife into three sections, which were homogenized in 1 mL of sucrose buffer (0.32 M sucrose, 10 mM HEPES, pH 7.4) freshly supplemented with protease inhibitors (Halt Protease Inhibitor Cocktail, 1861278, ThermoFisher) and WGA (100 μg/mL, L9640, Sigma-Aldrich). The homogenization consisted of 6–8 strokes at 500 rpm in a Teflon-glass tissue grinder, using an overhead stirrer (OS20L, Benchmark). The homogenate was centrifuged at 1200 × g for 10 min (4◦C) in a Legend Micro 21 high-speed centrifuge (ThermoFisher). After centrifugation, 850 μL of the supernatant 1 (S1) was transferred into a clean tube to obtain the P2 fraction; the remaining S1 fraction was removed to yield a pure precipitate 1 (P1) fraction, which was immediately fixed (see below). The tube containing the S1 fraction (850 μL) was centrifuged at 12,500 × g for 20 min (4◦C), and the resulting supernatant 2 (S2) was discarded using a vacuum. The pellet (precipitate 2, P2) was disaggregated by gentle agitation and by pipetting up-and-down 30 times in 1 mL of sucrose buffer. P2 protein concentration was determined by the BCA assay (ThermoFisher) using BSA standards.

The P1 nuclear fraction was fixed (Fix solution 1X; True-Nuclear Transcription Factor Buffer Set, Biolegend, #424401) for 10 min (light-protected) with rotation (RT), and centrifuged at 2000 × g for 5 min. Immediately after centrifugation, the supernatant was removed, and the pellet was resuspended in NIB buffer and washed twice with NIB by centrifugation at 2000 × g for 5 min (4◦C). After washing, the pellet was resuspended in 1 mL of NIB and was filtered using pre-wetted MACS® SmartStrainers (30 μm; Miltenyi, #130–098–458). Protein concentration was determined by the BCA assay (ThermoFisher) using BSA standards. A volume containing 100 μg of nuclear protein was transferred into cytometry tubes and washed with 2 mL of Perm/Wash Buffer 1X (True-Nuclear Transcription Factor Buffer Set) by centrifugation at 2000 × g for 5 min (20°C) using a swinging rotor (5910R centrifuge, Eppendorf). After centrifugation, the supernatant was discarded, and the pellet was resuspended before adding 250 µl of Perm/Wash Buffer with primary antibodies. Tubes were incubated for 1 h (RT) with shaking and washed with 2 mL of Perm/Wash Buffer, then centrifuged (2000 × g, 20°C, 5 min). Supernatant was discarded; the pellet was resuspended and incubated with an appropriate secondary antibody mixture in 250 µl of Perm/Wash Buffer for 30 min (RT) with shaking and in the dark. Finally, samples were washed with 4 mL of 0.1 % Triton-X and centrifuged (2000 × g, 20°C, 5 min). The supernatant was discarded, and the pellet was resuspended in 350 μL of PBS for flow cytometry analysis.

The P2 crude synaptosome fraction was fixed with 2% PFA for 15min (RT) with rotation. After incubation, 4 mL of NIB was added, and samples were centrifuged (4000 × g, 20°C, 10 min) using a swinging rotor (5910R centrifuge). The supernatant was discarded, and the pellet was resuspended in blocking buffer (0.32 M sucrose, 1mM EDTA, 5mM Tris, BSA 1.25% and FBS 1.25%). Permeabilization was done by incubating in 0.2% Tween 20 in blocking buffer for 15min (RT) with rotation. For immunostaining, 50μg of synaptosomes (quantified by BCA assay) were incubated (60 min, RT, with shaking) with primary antibodies in 250μL. Primary antibodies were washed with 4mL of 0.1% Tween 20 in SET buffer (0.32 M sucrose, 1mM EDTA, 5mM Tris) and centrifuged (4000 × g, 20°C, 10 min) in a swinging rotor (5910R centrifuge). Supernatant was discarded, and the pellet was resuspended in 250μL of appropriate secondary antibodies in 0.2% Tween 20 in blocking buffer and incubated (30min, RT) with shaking in the dark. Secondary antibodies were washed with 4mL of 0.1% Tween 20 in SET buffer and centrifuged (4000 × g, 20◦C, 10min). The pellet was finally resuspended in 500μL SET buffer for flow cytometry analysis.

Data were acquired using an Attune NxT flow cytometer (Thermo Fisher Scientific) equipped with acoustic focusing and blue (488nm) and red (638nm) lasers. Multiple fluorophores and dyes were detected by specific channels: Alexa Fluor 488 and EGFP with the BL1 channel (green, 530/30); PE and PI with the BL2 channel (orange, 574/26); StarBright Blue 765 and FM4–64 with the BL4 channel (near IR, 780/ 60); Alexa Fluor 647 with the RL1 channel (far red, 670/14); and CF770 with the RL3 channel (near IR, 780/60). For hierarchical gating on nuclear fractions, the ‘nuclei’ gate was defined based on the distribution of PI-positive events in the FSC-SSC plot, with the threshold set in FSC-H (125,000), and the stop criteria defined as 100,000 events or 250 µl. For P2 fractions, fluorescence triggering was set on BL4 (threshold = 500), and the stop criteria were defined as 30,000 events or 200 μl, with a flow rate of 200. We used FMOs as staining controls to define the threshold for positive signals; in particular, the FMO for EGFP was prepared with the dissected ventral hippocampus. We included a doublet discrimination analysis on BL4-H vs. BL4-A. Fluorescence spillover was corrected by compensation using single-stained samples, either P1- or P2-derived, to generate compensation matrices. Data analysis was performed using FlowJo V.11 software (LLC).

**Statistics**

Data are presented as mean ± s.e.m., box-and-whisker plots, and individual data points, as indicated in the figure legends. Sample sizes were determined based on preliminary experiments and previously published studies. Statistical analyses were performed using parametric tests, including two-tailed Student's *t*-tests and one-way, two-way, or repeated-measures ANOVA, as appropriate, when assumptions of normality and homogeneity of variance were met. Normality was assessed using the Kolmogorov–Smirnov test, and homogeneity of variance was assessed using Bartlett's test. When ANOVA indicated a significant effect, Šídák's multiple-comparisons test was used for all pairwise comparisons. When the assumptions for parametric testing were not met, appropriate nonparametric tests, including the Mann–Whitney U, Wilcoxon, or Kruskal–Wallis. Statistical analyses were performed using GraphPad Prism 9.0. All statistical tests were two-sided, and $P < 0.05$ was considered statistically significant.
